# Set1-dependent H3K4 methylation becomes critical for DNA replication fork progression in response to changes in S phase dynamics in *Saccharomyces cerevisiae*

**DOI:** 10.1101/2020.10.26.355008

**Authors:** Christophe de La Roche Saint-André, Vincent Géli

## Abstract

DNA replication is a highly regulated process that occurs in the context of chromatin structure and is sensitive to several histone post-translational modifications. In *Saccharomyces cerevisiae*, the histone methylase Set1 is responsible for the transcription-dependent deposition of H3K4 methylation (H3K4me) throughout the genome. Here we show that a combination of a hypomorphic replication mutation (*orc5-1*) with the absence of Set1 (*set1Δ*) compromises the progression through S phase, and this is associated with a large increase in DNA damage. The ensuing DNA damage checkpoint activation, in addition to that of the spindle assembly checkpoint, restricts the growth of *orc5-1 set1Δ*. Interestingly, *orc5-1 set1Δ* is sensitive to the lack of RNase H activity while a reduction of histone levels is able to counterbalance the loss of Set1. We propose that the recently described Set1-dependent mitigation of transcription-replication conflicts becomes critical for growth when the replication forks accelerate due to decreased origin firing in the *orc5-1* background. Furthermore, we show that an increase of reactive oxygen species (ROS) levels, likely a consequence of the elevated DNA damage, is partly responsible for the lethality in *orc5-1 set1Δ*.

**Author summary:** DNA replication, that ensures the duplication of the genetic material, starts at discrete sites, termed origins, before proceeding at replication forks whose progression is carefully controlled in order to avoid conflicts with the transcription of genes. In eukaryotes, DNA replication occurs in the context of chromatin, a structure in which DNA is wrapped around proteins, called histones, that are subjected to various chemical modifications. Among them, the methylation of the lysine 4 of histone H3 (H3K4) is carried out by Set1 in *Saccharomyces cerevisiae*, specifically at transcribed genes. We report that, when the replication fork accelerates in response to a reduction of active origins, the absence of Set1 leads to accumulation of DNA damage. Because H3K4 methylation was recently shown to slow down replication at transcribed genes, we propose that the Set1-dependent becomes crucial to limit the occurrence of conflicts between replication and transcription caused by replication fork acceleration. In agreement with this model, stabilization of transcription-dependent structures or reduction histone levels, to limit replication fork velocity, respectively exacerbates or moderates the effect of Set1 loss. Last, but not least, we show that the oxidative stress associated to DNA damage is partly responsible for cell lethality.

## Introduction

Replication of chromosomal DNA is a mechanism conserved among eukaryotes that is central for faithful genome propagation across cell divisions. It initiates at discrete genomic loci called origins of replication that are distributed along chromosomes [1]. The origins of replication in *Saccharomyces cerevisiae* are defined by short DNA sequences called autonomous replicating sequences (ARS) that are the sites of the cell cycle regulated assembly of prereplicative complexes (pre-RCs). The origin recognition complex (ORC) binds to ARS and, together with Cdc6 and Cdt1, is required for the loading of the Mcm2-7 complex, the replicative DNA helicase, during the G1-phase of the cell cycle. S-phase entry involves activation of the pre-RCs through phosphorylation of Mcm2-7 and recruitment of additional factors, converting the pre-RC into a pair of diverging replisomes [1]. The progression of the replication fork relies on the activated form of the helicase known as Cdc45/Mcm2-7/GINS (CGM) complex, which unwinds the DNA, followed by the DNA polymerases that carry out the synthesis of DNA at the leading and lagging strands [1].

Among the various obstacles that can be encountered by replication forks during their progression, those linked to transcription are potentially a major challenge [2]. Transcription-replication conflicts (TRCs) are prominent when the transcription machinery and the replisome are converging (head-on orientation). TCRs can result from the presence of either the transcription machinery itself or some transcription-induced/stabilized non-B DNA structures, such as R-loops which consist of DNA-RNA hybrids together with displaced ssDNA [2]. If sufficiently stable, R-loops can lead to the stalling of forks which, due to polymerase/helicase uncoupling and/or nucleolytic resection, are also a source of ssDNA [3]. The ssDNA of R-loops and stalled forks is rapidly coated by the ssDNA binding protein RPA [4,5]. The accumulation of RPA-coated ssDNA provides the signal for the recruitment of the Mec1/ATR checkpoint kinase to the defective replication fork [6–7]. The activation of Mec1 which results from this interaction can block mitosis in response to replication defects [8]. The processing of R-loops and stalled forks, that may eventually collapse, lead to the formation of double-strand breaks (DSBs). In this case, the repair of DSBs and therefore the possible restart of the fork depends on Rad52-dependent homologous recombination (HR) to repair the broken DNA[9,10].

As all DNA-templated processes, DNA replication is influenced by chromatin organization and its dynamics. Local chromatin structure, through nucleosome positioning, influences origin selection and function. The ARS sites bound by ORC are typically found within a nucleosome-free region (NFR) flanked by positioned nucleosomes on either side [11]. Although nucleosomes can interfere with the ORC DNA binding capacity, the interaction of ORC with ARS-adjacent nucleosomes [12] is thought to enforce origin specificity by suppressing non-specific ORC binding [13]. The compaction of DNA into chromatin poses a significant challenge to replication fork progression which must disrupt nucleosomes in order to progress [13,14]. At the same time, chromatin must be reconstituted behind the replication fork. Several chromatin remodeling enzymes have been reported to play important roles during DNA replication [13,15]. In addition to chromatin remodellers, various histone post-translational modifications are involved in DNA replication. For example, the acetylation of histones H3 and H4 stimulates the efficiency and accelerates the timing of origins firing within S phase [16,17], and the histone H3 lysine 56 acetylation by Rtt109 is a replication-associated mark that cooperates with the histone chaperone Asf1 to maintain a normal chromatin structure and is critical for cell survival in the presence of DNA damage during S phase [18,19]. The replication-coupled nucleosome assembly process can be influenced by histone modifications, as illustrated by the involvement of the histone lysine acetyltransferase Gcn5 in the deposition of histone H3 onto replicating DNA [20] and its contribution to maximum DNA synthesis rates *in vitro* [13]. H2B ubiquitylation promotes the assembly or stability of nucleosomes on newly replicated DNA, and this function is postulated to contribute to fork progression and replisome stability [21].

Some functional links between H3K4 methylation and replication have been described. In yeast, plants and mammalian cells, H3K4me2 and H3K4me3 are enriched on replication origins [22–24]. In human cells, the near-universal presence of dimethylated H3K4 at ORC2 sites led to consider it as a candidate mark recognized by ORC [25], and H3K4me3 demethylation was shown to promote replication origin activation by driving the chromatin binding of Cdc45 [26]. Additional work provided evidence that the MLL complexes and methylated H3K4 are involved in DNA replication through the replication licensing process [27]. Altogether, these studies points to a role of H3K4 methylation in DNA replication at the level of origin function. In *Saccharomyces cerevisiae*, the Set1-dependent H3K4 dimethylation has been described to promote replication origin function [22]. This would partly explain why *set1Δ* cells display a delayed entry into S phase in mitotic cells [28], as well as meiotic cells [29], although in mitotic cells this delay was proposed to be caused by Set1 contribution to proper cell-cycle-dependent gene expression [28]. Furthermore, Set1-dependent H3K4 dimethylation requirement for efficient origin activation [22] was not supported by an independent study where the recruitment and activation of DNA replication initiation factors was unaffected by the loss of H2B ubiquitylation [21], a context in which H3K4 dimethylation is absent [30]. Such discrepancy could be related to the fact that distinct replication origins were analyzed with alternative experimental methods. However, it is also possible that H3K4 methylation could influence DNA replication independently of origin firing as in the case of H2B ubiquitylation [21]. Recent works described the involvement of Set1 in DNA replication beyond origin firing. First, Set1 is important for the completion of DNA replication after an acute exposure to hydroxyurea (HU) [31]. The slower recovery of *set1Δ* from HU exposure is not the consequence of a defect in origin initiation but most likely in the processing of HU-stalled forks. Second, Set1 has a role in protecting the genome from TRCs occurrence with the transcription-deposited H3K4 methylation decelerating the fork progression at highly transcribed regions [32].

We have reinvestigated the conjecture that the genetic interactions that exist between *set1Δ* and replication-initiation mutants are related to origin firing defects [22]. We found that, when associated to the *orc5-1* mutation, that affect the subunit 5 of the ORC complex, the loss of Set1 leads to a strong defect of S-phase progression in mitotic and in meiotic cells. We failed to correlate this to a defect in origin function in *orc5-1 set1Δ* and instead we observed a strong increase in both the number and intensity of nuclear RPA and Rad52 foci, a sign of DNA damage accumulation in the double mutant. Accordingly, the DNA damage checkpoint acts together with the spindle assembly checkpoint to restrict the growth of *orc5-1 set1Δ* cells. Because *orc5-1 set1Δ* viability is decreased by the lack of RNase H activity but rescued by the reduction of histone gene dosage, we propose that the role Set1 plays in TRCs limitation [32] becomes critical when replication fork velocity is increased due to the *orc5-1* mutation. On another hand, *orc5-1 set1Δ* cell death is partly the consequence of reactive oxygen species (ROS) accumulation, likely in response to DNA damage.

## Results

### Cell proliferation and viability of *orc5-1* is promoted by Set1

We first confirmed, in the W303 background, the existence negative genetic interactions between the *SET1* deletion and temperature-sensitive alleles of various replication factors involved in origin function [22] : *orc5-1* (Fig 1), *cdc6-4* and *mcm2-1* (S1 Fig). Such an interaction was not seen with *cdc17-1* (a thermosensitive allele of DNA polymerase I alpha) (S1 Fig) which affects a step downstream of origin firing.

**Fig 1.**
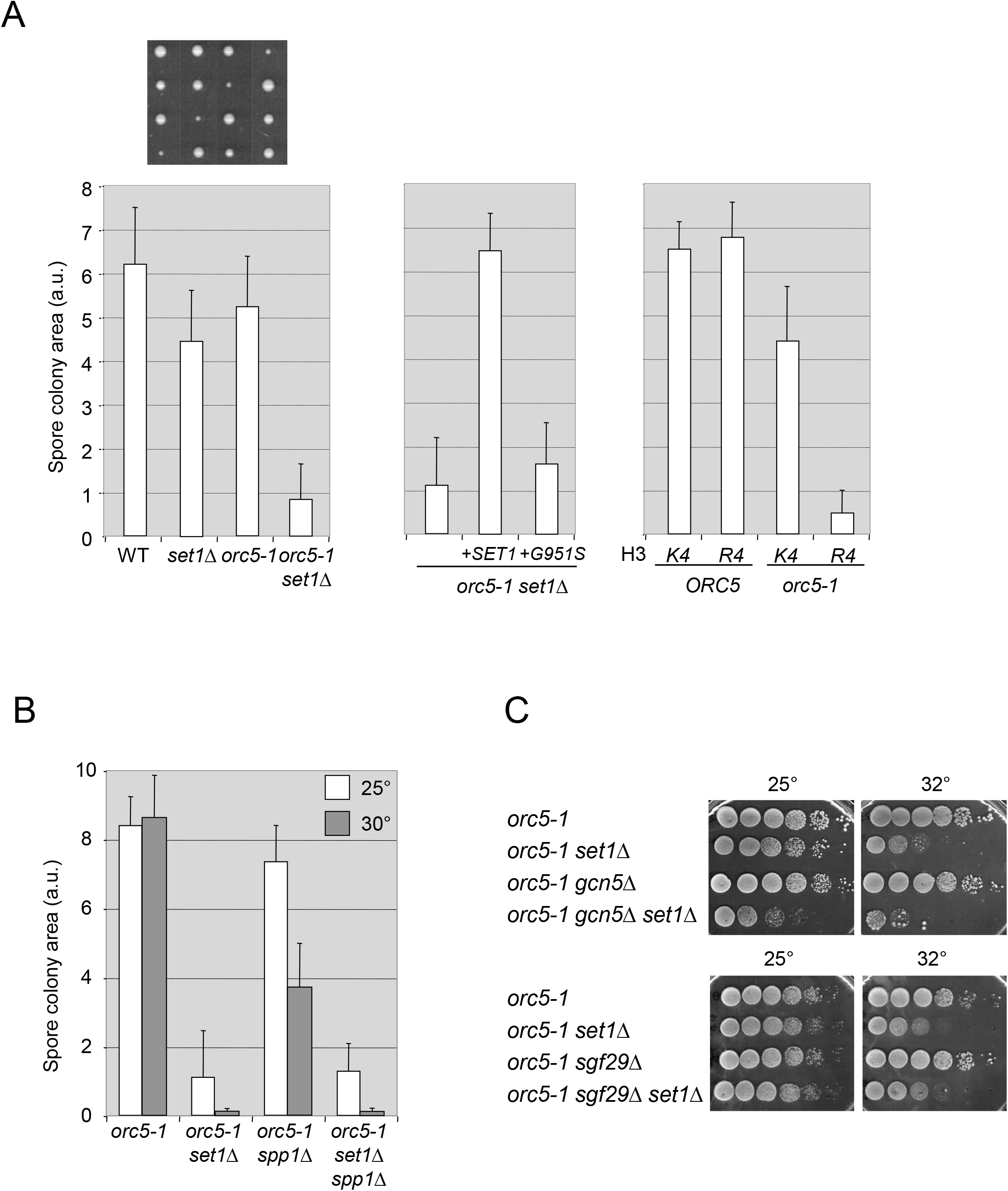
Negative genetic interaction between *set1Δ* and *orc5-1*. (A) Left: The deletion of *SET1* results in growth defect when combined with *orc5-1*. Growth of spore colonies after tetrad dissection of the diploid strain *set1Δ/SET1 orc5-1/ORC5*. The area of single-spore derived colonies was determined after three days at 25° since spore isolation (a.u.: arbitrary units). Error bars represent standard deviations of n≥15 spore colony area per genotype. Four representative tetrads (vertical lines), each with one small *orc5-1 set1Δ* colony, are shown on the top. Middle: Colony growth from *orc5-1 set1Δ* spores with an additional copy of *SET1* encoding either a wild-type (+*SET1)* or a catalytically inactive (+*G951S*) Set1 protein. Right: Colony growth from *ORC5* and *orc5-1* spores with wild-type copies (*K4*) and *K4R*-mutated copies (*R4*) of the H3 encoding genes *HHT1* and *HHT2*. (B) Effect of *SPP1* deletion is minimal compared to that of *SET1*. Growth of spore colonies, at 25° and 30°, after tetrad dissection of the diploid strain *set1Δ/SET1 spp1Δ/SPP1 orc5-1/ORC5*. (C) Thermosensitivity of *orc5-1 set1Δ* is independent of H3K4 acetylation. Tenfold serial dilutions of the respective strains were spotted onto YPD plates and incubated at the indicated temperatures for 3-4 days.

We focused on the *orc5-1* mutation as the genetic interaction with *set1Δ* was evident at 25° (see below), knowing that even at 23° only a subset of origins are activated in *orc5-1* [33]. Meiotic tetrads from a heterozygous *set1Δ/SET1 orc5-1*/*ORC5* diploid were dissected and the isolated spore colonies were grown at 25°. The colonies corresponding to *orc5-1 set1Δ* spores grew much more slowly than those of each single mutant (Fig 1A, left). This growth defect was exacerbated at 30° (Fig 1B), and at 32° the viability of *orc5-1 set1Δ* was severely compromised (Fig 1C).

To demonstrate that the slow growth phenotype of *orc5-1 set1Δ* was indeed caused by the lack of *SET1*, we performed a rescue experiment by adding a wild-type copy of *SET1* on a different chromosome (Fig 1A, middle). The complete restoration of the colony growth ruled out a possibility that *SET1* deletion might have disturbed expression of the adjacent *ORC6* gene thus causing an indirect effect. Notably, the *set1-G951S* allele encoding a catalytically dead version of Set1 was not able to rescue the growth of the double mutant, indicating that the Set1 histone methylase activity is required. Furthermore, the impact of *set1Δ* was recapitulated when the two copies of histone H3 bear the K4R (unmethylatable) mutation (Fig 1A, right), confirming the importance of H3K4 methylation for robust growth of the *orc5-1* mutant.

The loss of the COMPASS complex subunit Spp1 is associated with a specific decrease of H3K4me3 levels, without affecting mono- and dimethylation of H3K4 [34]. Compared to *set1Δ*, the negative impact of *spp1Δ* on the growth of *orc5-1* was moderate (Fig 1B), indicating that H3K4 trimethylation is partly dispensable for the growth of *orc5-1*, in contrast to mono- or dimethylation. Alternatively, this difference could be related to the larger increase of H3K4 acetylation levels in *set1Δ* compared to *spp1Δ* [35]. H3K4 acetylation mainly occurs in the absence of H3K4 methylation and essentially depends upon Gcn5, the catalytic subunit of different histone acetyltransferase (HAT) complexes, which is involved in the regulation of origin firing [16]. We therefore tested the impact of impairing H3K4 acetylation, through *GCN5* inactivation, on the thermosensitivity of *orc5-1.* Whereas the loss of Gcn5 had no effect by itself on *orc5-1* growth at 32°, the combination of *set1Δ* with *gcn5Δ* strongly affected the growth of *orc5-1* even at 25° (Fig 1C). Sgf29 is another component of the Gcn5-dependent HAT complexes that is involved in their recruitment to chromatin, through its interaction with H3K4me2/3 [36]. The negative effect of *set1Δ* on *orc5-1* growth at 32° was similar in the absence of Sgf29 (Fig 1C). Altogether, these results suggest that it is the lack of H3K4 methylation rather than the concomitant increase of Gcn5-dependent H3K4 acetylation that is responsible for the negative effect of *set1Δ* on *orc5-1* growth.

### The primary cause of the *orc5-1 set1Δ* genetic interaction is not related to a defect in origin activation

Since replication origin function is affected in *orc5-1* [33] and the involvement of H3K4 methylation in the same process has been described [22], an additive defect at the level of origin activity could be the basis of the *orc5-1 set1Δ* genetic interaction. The conclusion of [22] was based on a plasmid loss assay that measures the ability of cells to maintain a minichromosome bearing a single replication origin and a centromere. However, according to a recent genome wide analysis, no replication initiation defects were detected in *set1Δ*, with the bulk of early origins being as efficiently activated as in the wild-type [31]. This global analysis however does not exclude some variation in the activity of individual origins. In any case, whether that defective origin function was in fact responsible for the observed Set1-dependent genetic interactions has not been thoroughly investigated [22].

Therefore, we set out to employ the plasmid loss assay to test whether the *orc5-1 set1Δ* genetic interaction is a manifestation of a synergistic defect in origin function. Both the rate of plasmid loss that we have measured in the wild-type and its increase in *set1Δ* (Fig 2, left) were in agreement with the previous work [22]. A similar increment was observed in *orc5-1 set1Δ* relative to *orc5-1,* while the latter had much higher rate of plasmid loss compared to wild type. Importantly, in the context of *orc5-1*, the increased rate of plasmid loss due to lack of Set1 appears marginal compared to the drastic drop in viability (Fig 1C). To exclude the possibility that this limited increase in plasmid loss was due to the saturation of the assay, the temperature of 25° was used in order to limit inactivation of the *orc5-1* encoded protein (Fig 2, right). Again, despite the milder rate of plasmid loss in *orc5-1*, no substantial increase was associated with the loss of Set1, contrasting with a strong genetic interaction observed for the growth rate (Fig 1A). While we cannot exclude some contribution, the activation of origins is likely not a determining factor in the *orc5-1 set1Δ* genetic interaction.

**Fig 2.**
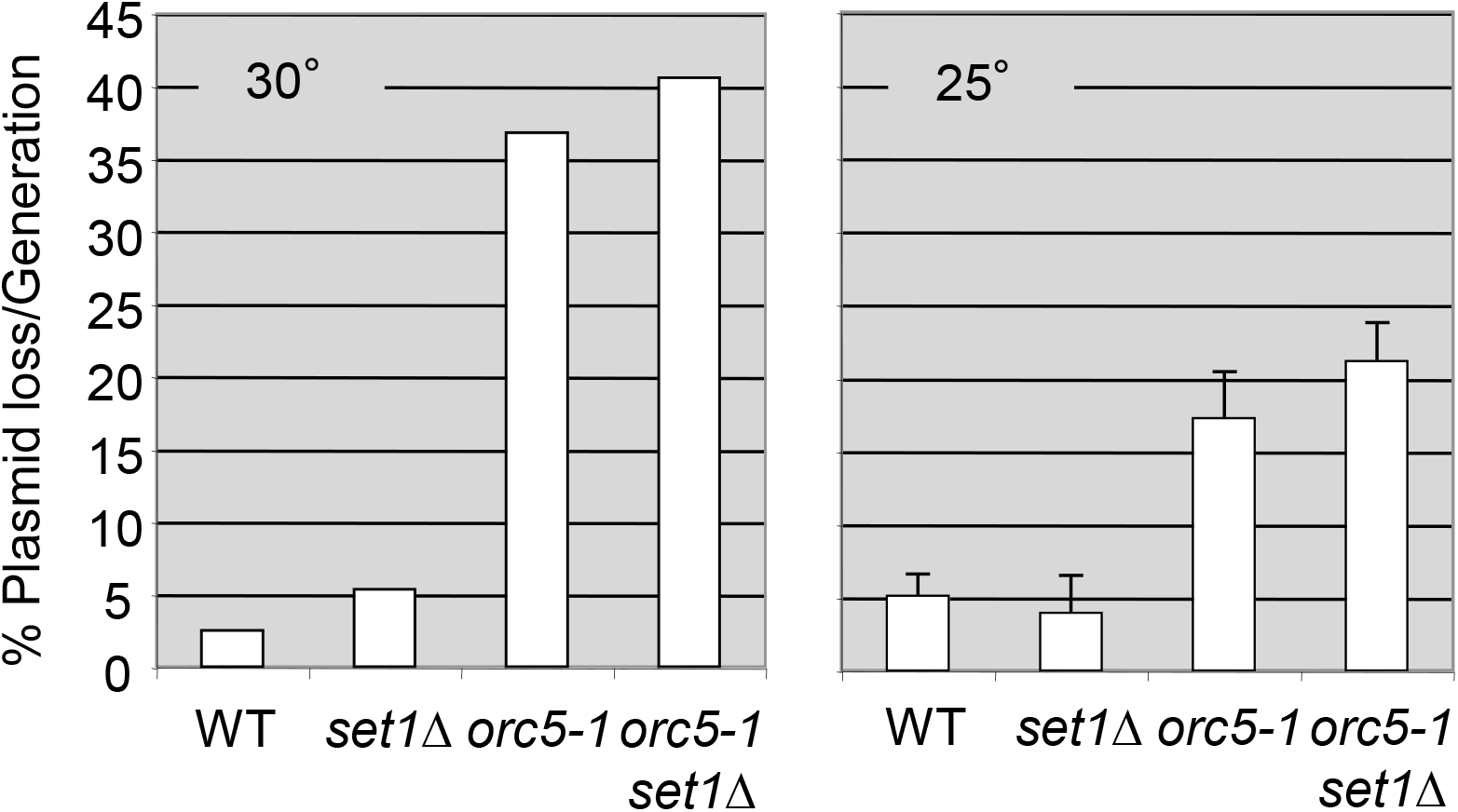
Testing the *orc5-1 set1Δ* genetic interaction through an origin-dependent plasmid maintenance assay. Plasmid loss rates of an ARS-CEN-bearing plasmid (pRM102) were measured in wild-type, *set1Δ, orc5-1* and *orc5-1 set1Δ* strains at the temperatures of 30° and 25°. Loss rates are reported per cell division (see Materials and Methods). For experiments at 25°, the average loss rates were obtained from three independent transformants for each strain, and the error bars indicate standard deviations.

### Progression through S phase is compromised in *orc5-1 set1Δ* cells

Analysis of the cell cycle profiles by flow cytometry did not reveal any significant changes in the distribution along the cell cycle in *orc5-1 set1Δ* during exponential growth at 25° (see −3h on Fig 3A). Cells were then arrested in late G1-phase by ⍰-factor treatment during two hours, shifted to 37° for an additional hour to inactivate *orc5-1*, and then released from the G1 block at 37° (Fig 3A). No delay in the progression through S phase was apparent for *orc5-1* cells that completed DNA replication after 30 min, similarly to the wild-type. This suggests that any decrease in origin firing, as evidenced by the plasmid loss assay (Fig 2), is compensated by an increase of the replication fork rate as shown previously for other mutations affecting the level of origin firing [37]. While *set1Δ* cells display a slight replication delay at 37°, the progression through S phase was notably slower in *orc5-1 set1Δ* with most of the cells having less than 2C DNA content when replication is complete in both single mutants. Therefore, Set1 loss appeared to affect S-phase progression when ORC function is compromised.

**Fig 3.**
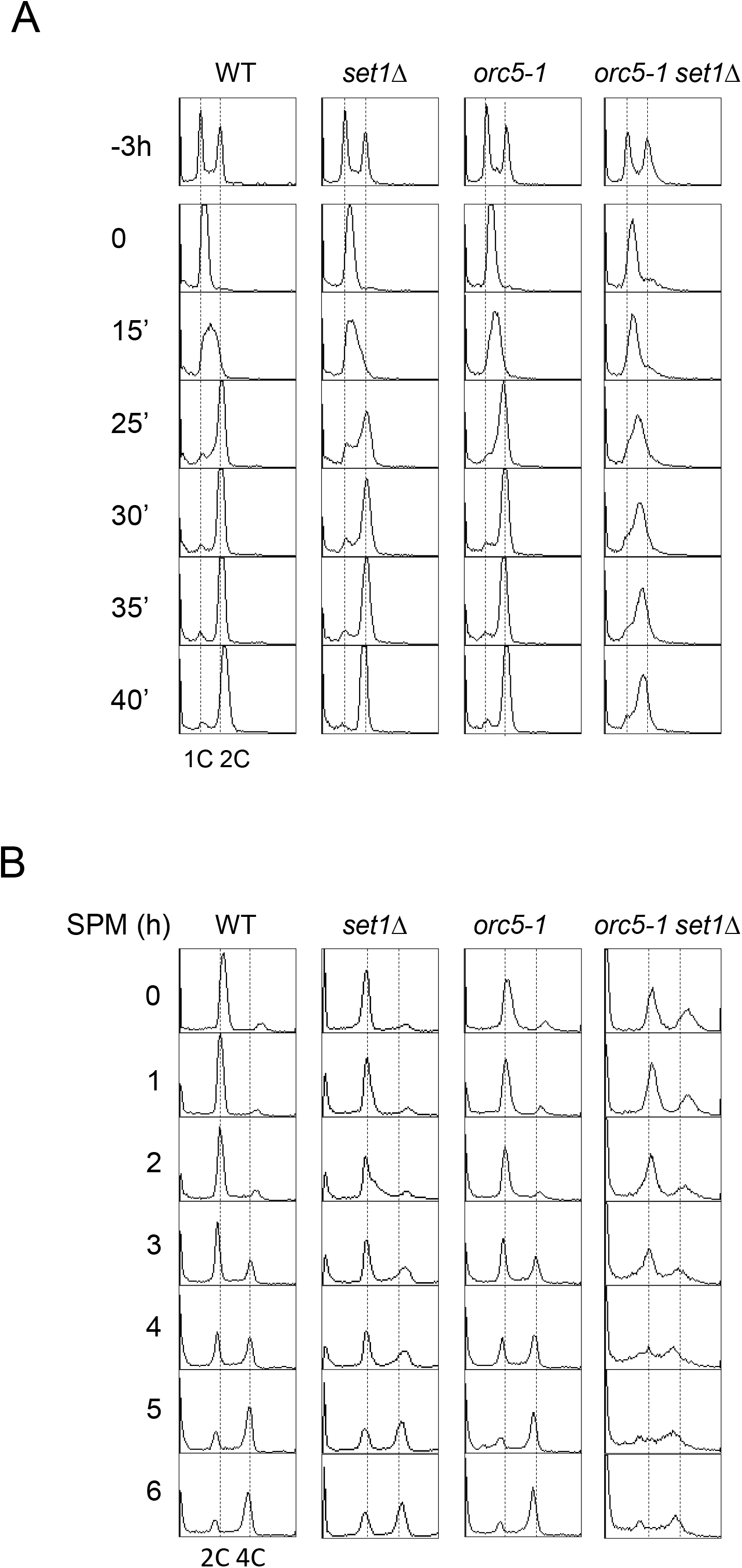
The progression through S phase is compromised in *orc5-1 set1Δ*. (A) Strains (W303 background) grown at 25° were treated with ⍰-factor during two hours followed by one hour at 37° before their release from the G1 block into fresh medium at 37°. DNA content was determined before (time −3h) and after synchronization (time 0), then at the indicated intervals (minutes). (B) After synchronization in YPA medium at 25° (time 0), cells were transferred at 30° in sporulation medium (SPM). Meiotic DNA replication of WT, *orc5-1*, *set1*Δ and *orc5-1 set1*Δ diploids (SK1 background) was monitored by FACS analysis at the indicated intervals (hours).

A progressive accumulation of cells with a 2C DNA content, suggesting a delay/arrest in G2/M, has been described for *orc5-1* cultures at non permissive temperature [39]. To determine whether the lack of Set1 would influence this tendency, the cell cycle profiles of non-synchronized *orc5-1* and *orc5-1 set1Δ* cultures were monitored for extended period of time after shift from 25° to 37° (S2 Fig). We found that unlike *orc5-1*, most of the *orc5-1 set1Δ* cells accumulated in S phase. In agreement with the results obtained with synchronized cultures (Fig 3), it appears that the progression through S phase becomes the main limiting step in *orc5-1 set1Δ* at 37°.

Finally, we tested whether the *orc5-1 set1Δ* genetic interaction extends to DNA replication that occurs during meiosis, i.e. in the conditions for which a clear impact of *set1Δ* has been described [29]. After successive backcrosses, the *orc5-1* mutation was introgressed into the SK1 background, which allows rapid and synchronous sporulation. Suitable diploids were further obtained. As a first indication of the genetic interaction between *orc5-1* and *set1Δ*, at 25° the sporulation levels of the *orc5-1 set1Δ* diploid was severely reduced compared to that of each single mutant (S3 Fig). After shifting G1-synchronized cells from 25° to 30°, the meiotic replication of SK1 diploids were compared (Fig 3B). As already published [29], the meiotic replication in *set1Δ* cells is delayed. In *orc5-1 set1Δ* cells, the decrease of the 2C peak, which reflects the entry into S phase, was not accompanied by a parallel increase of the 4C peak and most of the cells appear eventually blocked throughout S phase. This correlates with the fact that only a fraction of *orc5-1 set1Δ* cells complete meiosis up to the stage of spore formation (S3 Fig). Thus, as in vegetative conditions, the progression through S phase in meiotic cells appears to be strongly affected when *set1Δ* is combined with the *orc5-1* mutation.

### Differential sensitivity of *orc5-1* and *orc5-1 set1Δ* to checkpoint inhibition

The involvement of the spindle assembly checkpoint (SAC) in response to defects in ORC function has been revealed by the mitigation of *orc1-161* associated cell lethality by the deletion of *MAD2* (Gibson et al., 2006). Similarly, we found that *mad2Δ* increased the viability of *orc5-1* at the non-permissive temperature of 35,5° (Fig 4A and S4A Fig for independent clones), suggesting that SAC activation restrains the growth of *orc5-1*. Although the way SAC is activated when ORC function is compromised is unclear, this could be linked to the contribution of ORC function to sister chromatid cohesion [40,41]. Such a rescue is not seen when the mitotic exit network regulator Bub2 is absent or in the presence of *rad53K227A*, a kinase deficient allele of *RAD53* (Fig 4A and S4A Fig). The fact that the *rad53-K227A* mutation aggravates the thermosensitivity of *orc5-1* even when Mad2 is missing could indicate that the functionality and/or structure of replication forks are compromised (see discussion). We further observed that the loss of Mad2 does not rescue the thermosensitivity of *orc5-1 set1Δ* at 35.5°, although a positive effect can be observed at the lower temperature of 34° (S4B Fig). This suggests that above a certain temperature, i.e. some degree of Orc5 inactivation, an additional Mad2-independent checkpoint is activated, which restrains proliferation in *orc5-1 set1Δ* even in the absence of SAC activity.

**Fig 4.**
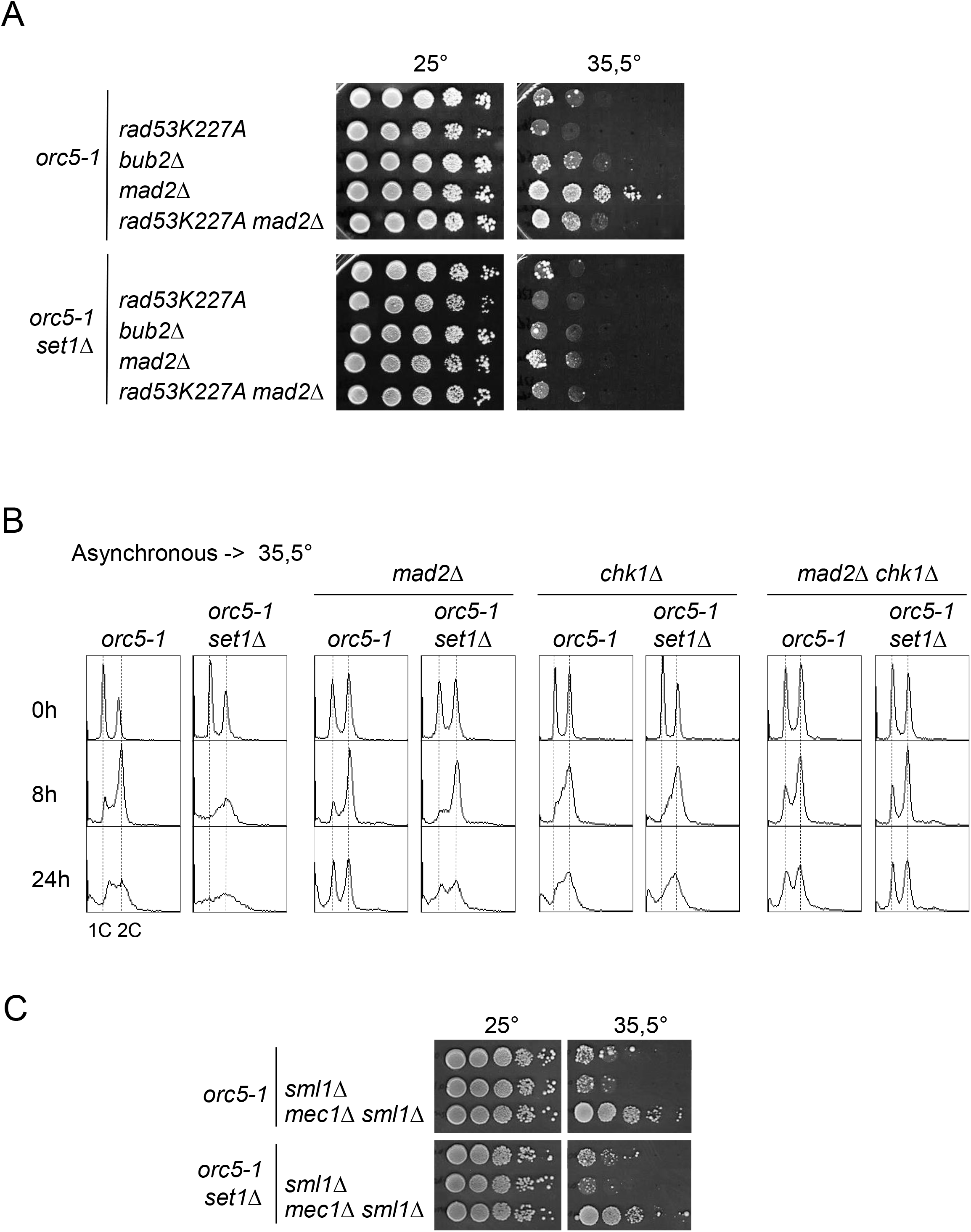
Distinguishable contribution of the spindle assembly checkpoint and the DNA damage checkpoint to cell cycle arrest in *orc5-1* and *orc5-1 set1Δ*. (A) Different impact of *mad2Δ* on the thermosensitivity of *orc5-1* and *orc5-1 set1Δ*. Tenfold serial dilutions of the respective strains were spotted onto YPD plates and incubated at the indicated temperatures for 3 (25°) or 4 (35,5°) days. (B) Unsynchronized cultures (25°) of the indicated strains were shifted to 35,5°. DNA content was determined at the indicated times (hours). 1C and 2C (broken lines) indicate the DNA content of cells with unreplicated or fully replicated DNA, respectively. (C) Rescue of *orc5-1* and *orc5-1 set1Δ* thermosensitivity by *mec1Δ*. Tenfold serial dilutions of the respective strains were spotted onto YPD plates and incubated at the indicated temperatures for 3 (25°) or 4 (35,5°) days.

This difference regarding the effect of Mad2 loss is reflected in the cell cycle profiles from liquid cultures, after shifting the cells to the same temperature of 35,5° (Fig 4B). In *orc5-1*, the absence of Mad2 has no impact on the short-term accumulation of cells with a 2C DNA content (8h) but is then followed by a normal cell cycle profile (24h) which is consistent with the rescue of colony growth observed on solid medium (Fig 4A). Such a return to a normal cell cycle is not observed for *orc5-1 set1Δ mad2Δ*, with a 24h cell cycle profile being similar to that of *orc5-1*, again in agreement with the absence of colony growth on solid medium. The transient G2/M accumulation observed when Mad2 is missing suggests that another pathway could limit the G2/M to anaphase transition. The critical target of the Mad2-dependent spindle checkpoint, the inhibitor of anaphase Pds1, is also targeted by the DNA damage checkpoint kinase Chk1[42]. Although with a limited effect on its own, the loss of Chk1 when Mad2 is absent strongly reduced the short-term G2/M accumulation (8h) and restored a normal cell cycle distribution in *orc5-1 set1Δ* after 24h (Fig 4B). Thus, whereas a defective Orc5 leads to SAC activation, the additional loss of Set1 appears to be associated with the activation of a DNA damage checkpoint that restrains *orc5-1 set1Δ* cell cycle progression in the absence of Mad2.

The major DNA damage checkpoint kinase Mec1 has been proposed to act upstream of both Mad2 and Chk1 in the control of the G2/M to anaphase transition [43]. According to this model, the inactivation of Mec1 would have the same effect as the simultaneous inactivation of Mad2 and Chk1. The essential role in cell viability of Mec1 can be bypassed by deletion of *SML1* which increases dNTP levels [44]. Although Sml1 inactivation has no positive effect on its own, excluding a role for limiting amounts of dNTPs, inactivation of Mec1 alleviated the cell multiplication restriction of *orc5-1* and *orc5-1 set1Δ* at 35,5° (Fig 4C). This supports the view that the DNA damage checkpoint limits the proliferation of *orc5-1* when Set1 is missing.

### The replication stress in *orc5-1 set1Δ* is associated to a specific increase of DNA damage

According to the results described above, the absence of Set1, when associated to *orc5-1*, should lead to a measurable increase of the signal that enables activation of the Mec1-Chk1 pathway, i.e. of the RPA-coated ssDNA. To address this issue, we analyzed foci of CFP fused to Rfa1, the largest subunit of RPA, as a readout of the amount of ssDNA present in the nucleus. The percentage of cells with Rfa1 foci was measured in log phase cells at 30°, a temperature permissive for *orc5-1*, to discern more easily the specific contribution of the Set1 loss on the amount of ssDNA detected in the *orc5-1 set1Δ*. Spontaneous Rfa1 foci were found in about 13% of WT cells (Fig 5A) the majority of which are observed during the G1 and S phases of the cell cycle, with some rare G2/M cells displaying the most intense foci (S5 Fig). While the percentage of cells with Rfa1 foci as well as the average focus intensity were similar in *set1Δ* and only slightly higher in *orc5-1* both were significantly increased in *orc5-1 set1Δ* (Fig 5A and S5 Fig) with a larger fraction of cells harboring more than one focus (see double arrow on microscopy photograph, Fig 5A). Thus, the impairment of fork progression as well as the DNA damage checkpoint activation in *orc5-1 set1Δ* is correlated to an excess of Rfa1 foci.

**Fig 5.**
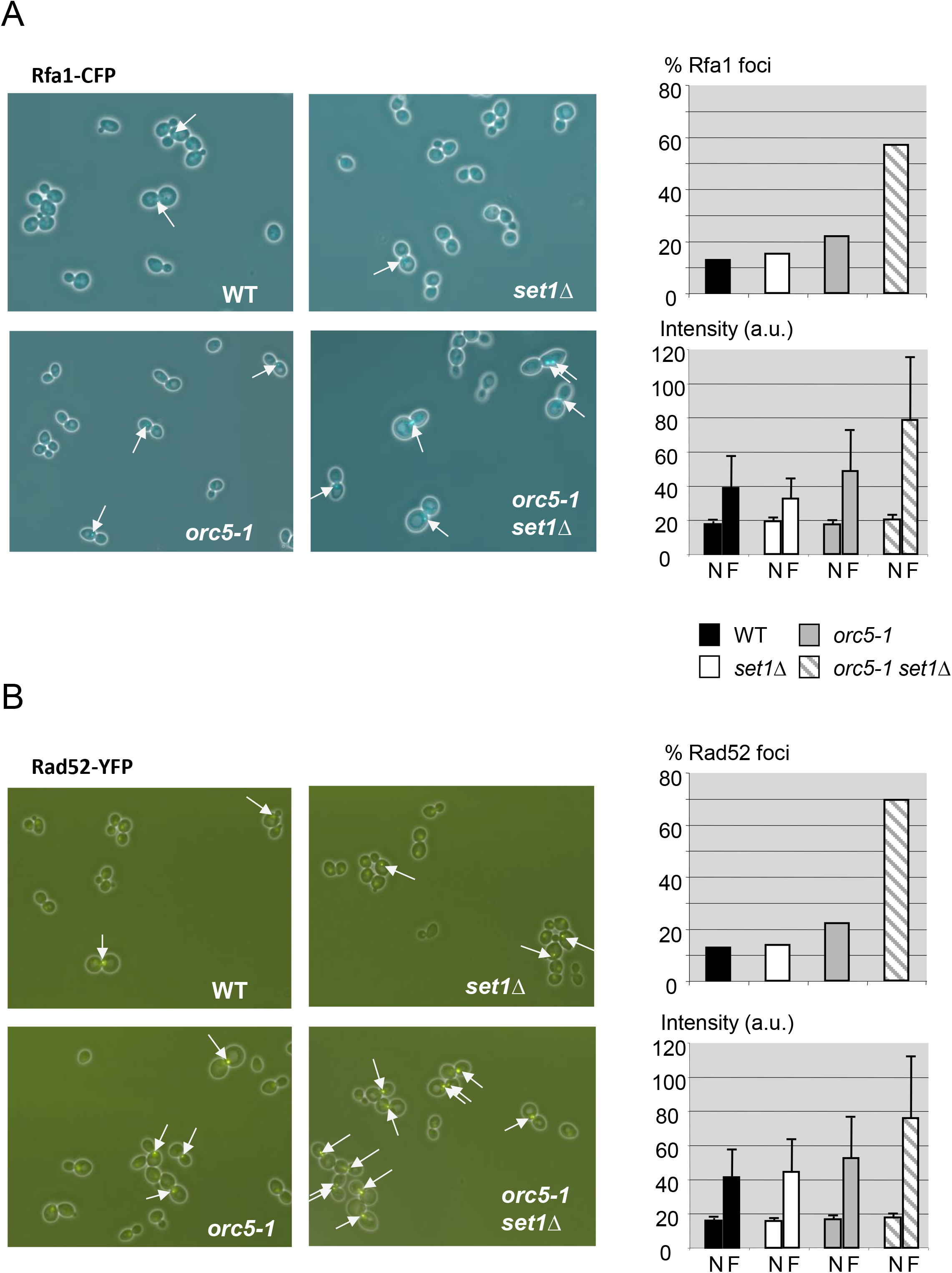
Elevated spontaneous DNA damage levels in *orc5-1 set1Δ* cells. (A) Left, microscopy photographs (merged CFP and differential interference contrast images) of cells expressing Rfa1-CFP. Arrows indicate foci. Top right, proportion (%) of cells containing Rfa1-CFP foci in exponentially growing cells (30°) of the indicated genotypes. Bottom right, average intensity (arbitrary units) of the Rfa1 foci (F) compared to the average nuclear background fluorescence (N). Error bars indicate standard deviations. (B) Left, microscopy photographs (merged YFP and differential interference contrast images) of cells expressing Rad52-YFP. Arrows indicate foci. Top right, proportion (%) of cells containing Rad52-YFP foci in exponentially growing cells (30°) of the indicated genotypes. Bottom right, average intensity (arbitrary units) of the Rad52 foci (F) compared to the average nuclear background fluorescence (N). Error bars indicate standard deviations.

Some RPA-coated ssDNA may correspond to resection products of DNA double-strand breaks (DSBs) associated to stalled fork processing, whose repair depends on the Rad52-dependent homologous recombination pathway [45]. Thus, we analyzed the presence of Rad52-repair centers by measuring Rad52 nuclear foci using a functional Rad52-YFP fusion protein. The results were very similar to those obtained with Rfa1, with more numerous and more intense Rad52 foci in *orc5-1 set1Δ* (Fig 5B and S5 Fig). As for Rfa1, the vast majority of the most intense Rad52 foci were present in *orc5-1 set1Δ* nuclei in agreement with the larger proportion of G2/M cells (S5 Fig). As a consequence, the survival of *orc5-1 set1Δ* cells must be particularly dependent on the Rad52-dependent DNA repair activity. We confirmed this by testing the impact of Rad52 loss on cell viability and proliferation (Fig 6). Although *orc5-1* was slightly more sensitive than the wild-type and *set1Δ* to the deletion of *RAD52* (S6 Fig), in agreement with a small increase in Rad52 foci (Fig 5B), the lack of Rad52 was clearly most deleterious in *orc5-1 set1Δ*. Thus, most *orc5-1 set1Δ rad52Δ* spores did not give a colony at 30° (Fig 6A) and when they did the rate of colony growth was severely affected at 25° (S6 Fig). Analysis of the DNA content of *orc5-1 set1Δ rad52Δ* at 30° shows an accumulation of cells with less than 1C DNA content, likely corresponding to dead cells (Fig 6B).

**Fig 6.**
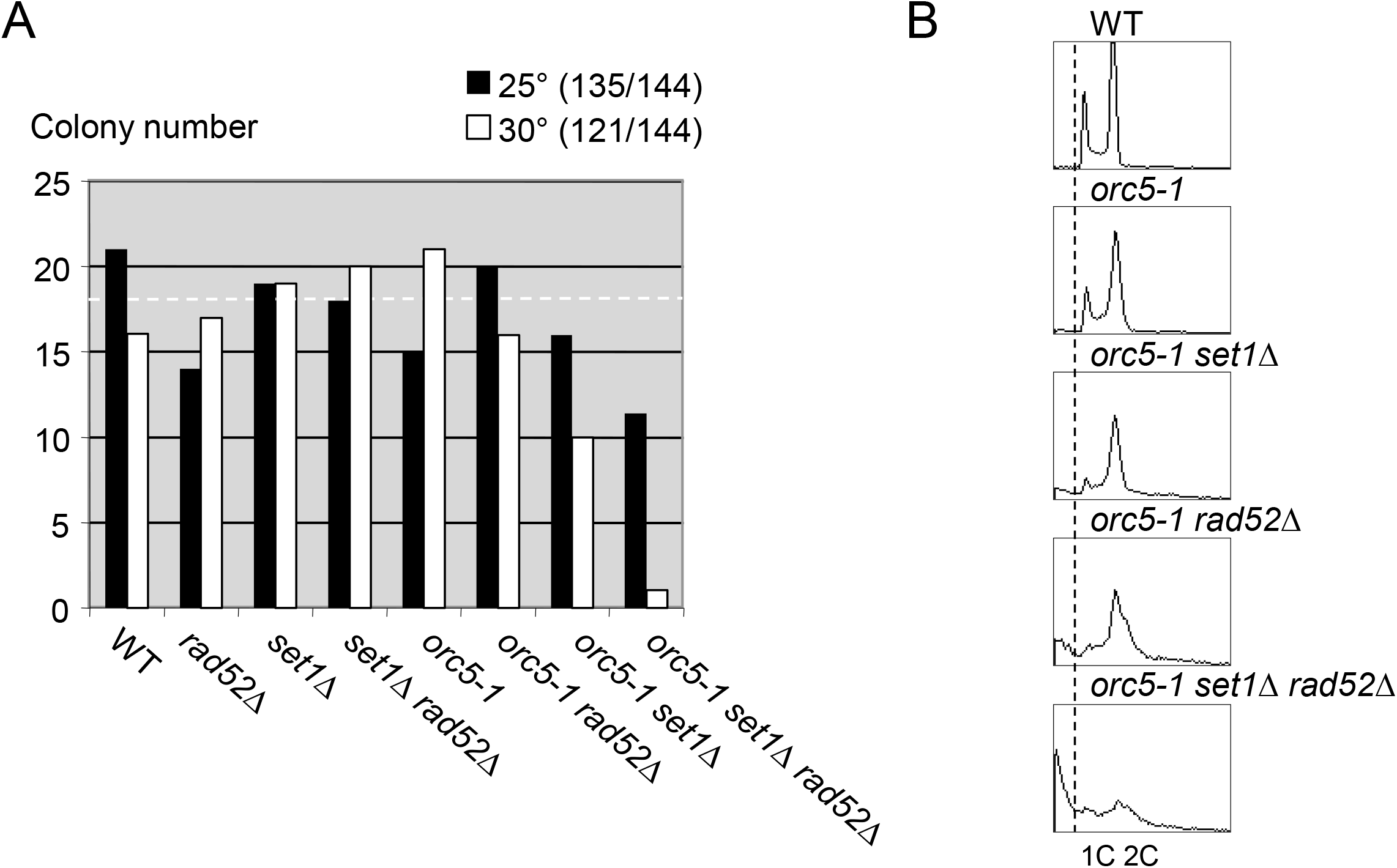
Viability of *orc5-1 set1Δ* cells depends on Rad52. (A) Numbers of spore colonies obtained for each genotype at 25° and 30° after tetrad dissection of the diploid strain *set1Δ/SET1 orc5-1/ORC5 rad52Δ/RAD52*. Brackets: total number of viable colonies / total number of isolated spores. The horizontal white broken line indicates the theoretical number of colonies (18 = 144 spores / 8 genotypes) expected for an even distribution of each genotype if all spores were viable. (B) Cells of the indicated genotype were cultured for 5 hours at 30° before their DNA content was analyzed by FACS. The fraction of cells with less than 1C DNA content (presumably apoptotic cells) is delimited by a vertical broken line.

The large increase of Rad52-dependent DNA repair activity associated with a slow progression though S phase unveils the replication stress that exists in *orc5-1 set1Δ*. Accordingly, one can expect a hypersensitivity of *orc5-1 set1Δ* cells to any form of additional stress that affect DNA replication. In fact, the viability of *orc5-1 set1Δ* was severely affected in the presence of hydroxyurea (HU), at concentrations sparing the viability of each single mutant (S7 Fig) indicating that lack of Set1 further aggravates fork progression in HU. Osmostress is an alternative way to impede replication fork progression [46]. Similarly, the viability of *orc5-1 set1Δ* was specifically affected when exposed to high osmolarity (S7 Fig).

### Transcription replication conflicts as a source of DNA damage in *orc5-1 set1Δ*

As RNase H is able to process co-transcriptional R-loops responsible for TRCs [47], we examined the effect of eliminating both RNase H1 (*rnh1Δ*) and RNase H2 (*rnh201Δ*) activities. The *orc5-1* cell viability at 31° is affected by the deletion of *RNH1* and *RNH201* (Fig 7A). One interpretation is that the increase of replication fork speed due to *orc5-1* makes them more prone to encounter R-loops stabilized by RNase H inactivation. The negative effects of *rnh1Δ rnh201Δ* are clearly stronger when Set1 is missing as shown by the greater sensitivity of *orc5-1 set1Δ* to the lack of RNase H activity. According to the recent description of the role of H3K4 methylation in decelerating ongoing replication at highly transcribed regions [32], the fact that replication fork rate increase is not counterbalanced by Set1 would lead to a substantial rise of TRCs severity in *orc5-1 set1Δ* as evidenced by the increase of Rad52 nuclear foci (Fig 5B). These results argue in favor of co-transcriptional R-loops as a source of DNA damage in *orc5-1 set1Δ*.

**Fig 7.**
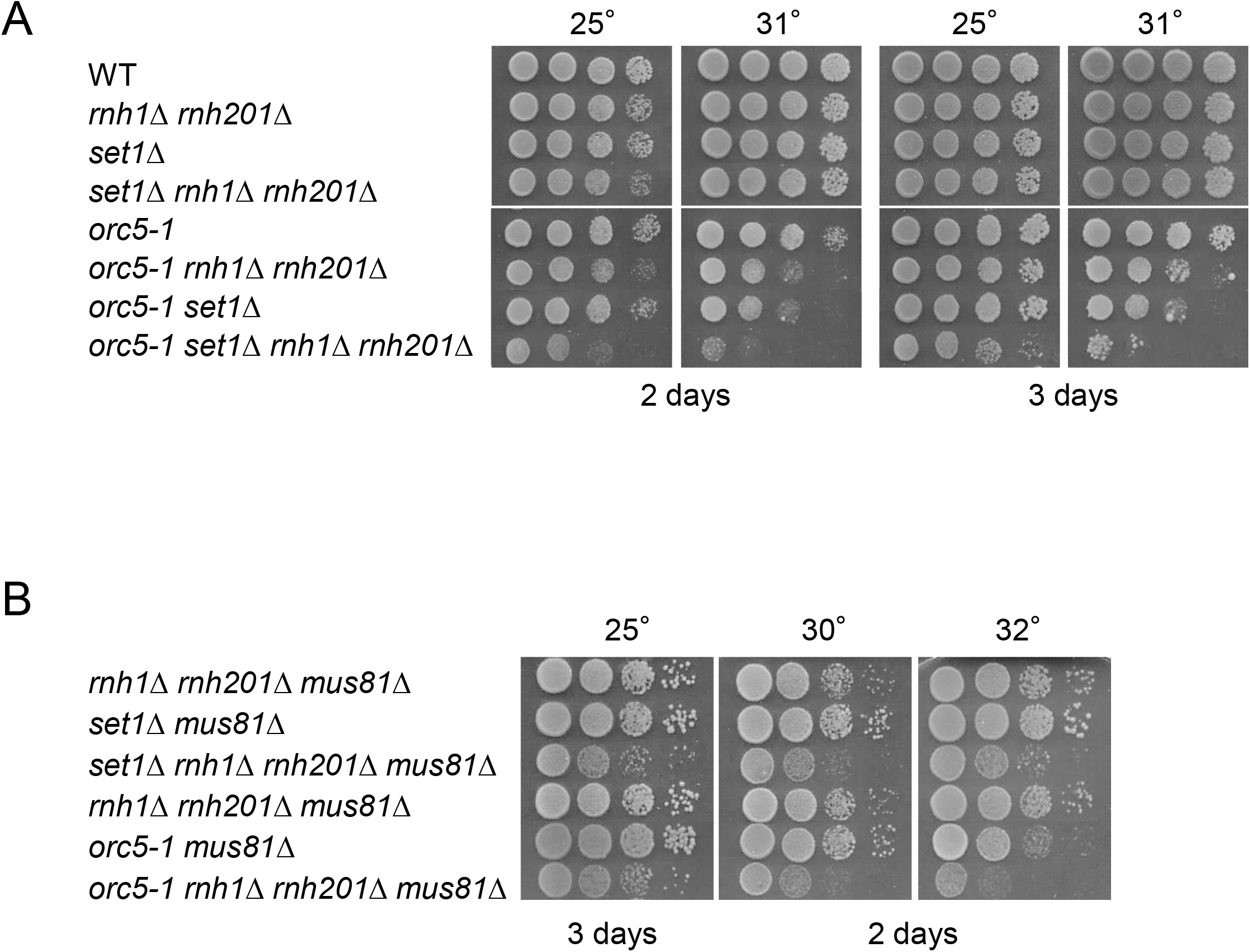
Loss of Set1 sensitizes cells to RNase H removal alone or together with Mus81. (A) and (B) Tenfold serial dilutions of the respective strains were spotted onto YPD plates and incubated at the indicated temperatures for 2 and 3 days.

Whereas Rad52 is required for any kind of DSBs, the Mus81-Mms4 resolvase is more specifically required for repair of one-ended DSBs formed at collapsed forks [47]. Moreover, Mus81 is involved in the protection and restart of forks stalled by co-transcriptional R-loops [48,49]. We found that, contrary to *Schizosaccharomyces pombe* [47], the *mus81Δ rnh1Δ rnh201Δ* combination is not lethal in *S. cerevisiae* (Fig 7B). However, the effects of the *mus81Δ* and the *rnh1Δ rnh201Δ* mutations are additive in *orc5-1* and, more strikingly, in *set1Δ*. Although the loss of Mus81 has little effect by itself (S8 Fig), this suggests that the lack of H3K4 methylation sensitize cells to the accumulation of R-loops when Mus81 is missing.

### Limiting histone supply improves the viability of *orc5-1 set1Δ*

As we hypothesized that the function of H3K4 methylation could become critical in *orc5-1* cells because of the compensatory increase of replication fork rate in *orc5-1* [37], decreasing replication fork velocity by other means is expected to attenuate the negative impact of Set1 loss. Because of the functional coupling between replication fork progression and nucleosome assembly, the fork speed can be restrained by decreasing histone levels [50]. In line with this, we found that deletion of *HHT2-HHF2*, one of the histone gene pairs encoding histone H3 and H4, completely suppressed the viability loss of *orc5-1 set1Δ* at 32° (Fig 8A). We also observed that *hht2-hhf2Δ* reduced the sensitivity to HU (50mM) of *orc5-1* and *orc5-1 set1Δ* cells (Fig 8B). Similar results were obtained by deleting *HHT1-HHF1,* the other histone gene pair (S9 Fig). A straightforward interpretation of these results is that limiting amounts of H3/H4, by mitigating the increase of fork rate due to *orc5-1*, can counteract the negative effect that *set1Δ* and HU have on *orc5-1* viability.

**Fig 8.**
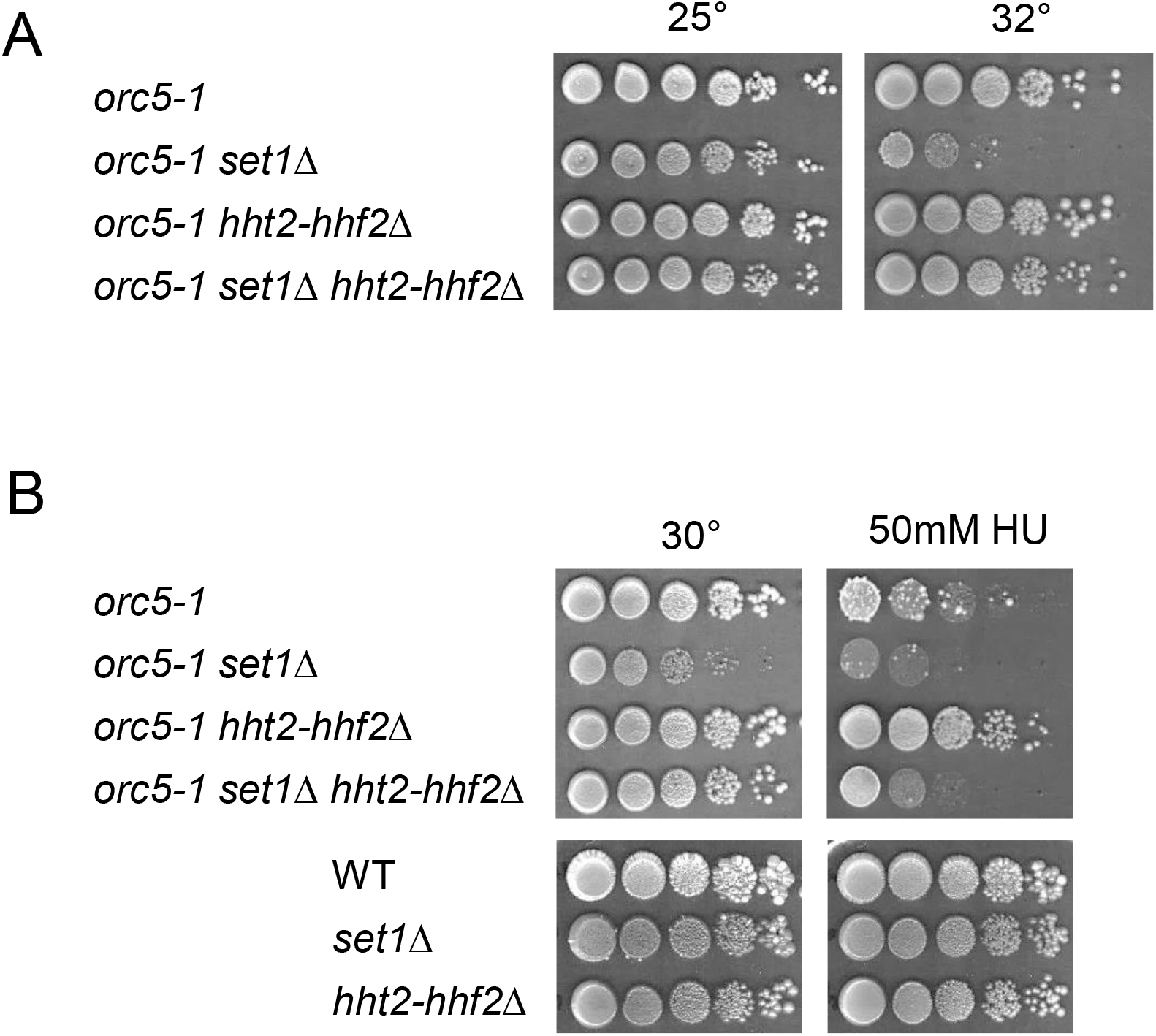
Reduction in histone gene dosage limits the negative effect of Set1 loss and hydroxyurea on *orc5-1*. (A) and (B) Tenfold serial dilutions of the respective strains were spotted onto YPD plates without or with HU (50mM) and incubated at the indicated temperatures for 3-4 days.

### Oxidative stress contributes to *orc5-1 set1Δ* lethality

Whereas DNA damage accumulation is most certainly responsible for the Mec1-dependent cell cycle arrest in *orc5-1 set1Δ*, whether it contributes directly to cell death is unknown. The elevation of reactive oxygen species (ROS) levels in *orc5-1* at the non-permissive temperature of 37° [51] and the fact that Set1 loss leads to ROS accumulation during aging [52], led us to consider the oxidative stress as a cause of lethality of *orc5-1 set1Δ*. Whereas intracellular ROS levels in the single mutants were similar to that in the wild type (at 25° and 32°), a clear increase was seen for *orc5-1 set1Δ* (Fig 9A). This increase correlated with the decrease in cell viability that was specifically observed for the double mutant at 32° (Figs 1C and 9B, top). The cell viability of *orc5-1 set1Δ* was improved in the presence of the antioxidant N-Acetylcysteine (NAC) (Fig 9B, top), whereas it was further aggravated by the ROS hydrogen peroxide (H_2_O_2_) (Fig 9B, middle). This indicates that ROS are, at least partly, responsible for the cell lethality of *orc5-1 set1Δ*. Such a causal link was reinforced by the mitigating effect the loss of the ROS responsive metacaspase Yca1 had on the *orc5-1 set1Δ* thermosensitivity (Fig 9B, bottom).

**Fig 9.**
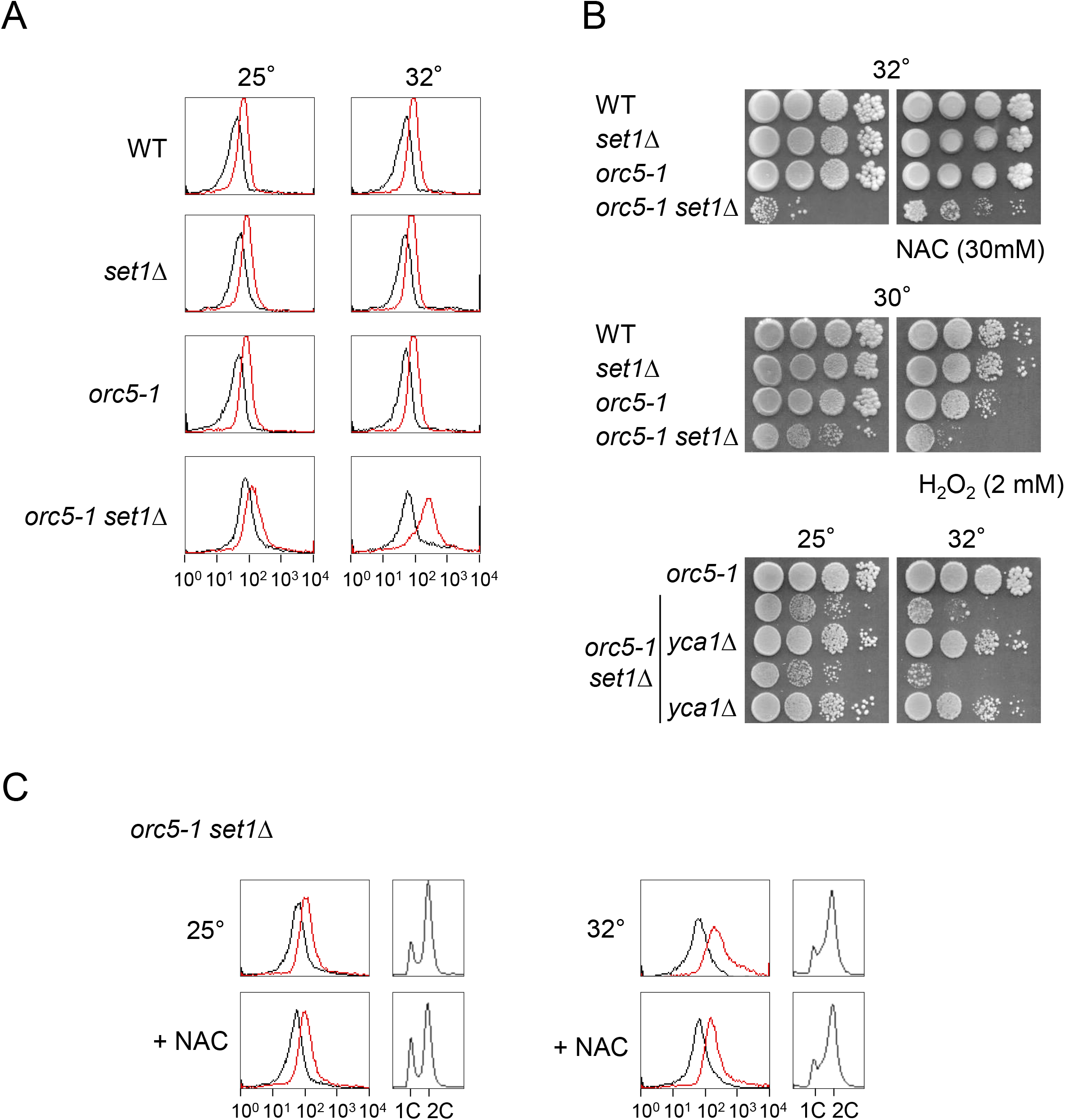
Oxidative stress is involved in *orc5-1 set1Δ* lethality. (A) The indicated strains were cultured at either 25°or 32° before ROS levels were determined at 7 hours. The fluorescence signals measured in absence (black) or presence (red) of the ROS-sensitive fluorochrome (CellROX green) are surperimposed. X-axis and y-axis: fluorescence intensity (log scale) and number of cells, respectively. (B) The viability of *orc5-1 set1Δ* is sensitive to NAC, H_2_O_2_ and *yca1Δ*. Tenfold serial dilutions of the respective strains were spotted onto YPD plates at the indicated temperatures for 3-4 days. NAC and H_2_O_2_ concentrations are indicated. (C) Decreasing ROS levels is without effect on the S-phase progression defect in *orc5-1 set1Δ*. Cells were cultured at 25°and 32°, in absence or presence of NAC (30mM), before ROS levels (left) and DNA content (right) were determined at 7 hours.

As both DNA damage and ROS levels are increased in *orc5-1 set1Δ*, the question arises whether a relationship exists between the two, with either ROS inducing DNA damage or the other way around. To get insight into this question, we tested whether ROS mitigation by NAC had an effect on the perturbed cell cycle distribution displayed by *orc5-1 set1Δ* at 32° (Fig 9C). For this purpose, *orc5-1 set1Δ* cells were grown at 25° and 32° in either presence or absence of NAC. At 25°, the addition of NAC had some discernible influence on the cell cycle profile, with an increase of the G1 to G2/M ratio. At 32°, the relative accumulation of cells along the S phase was insensitive to the presence of NAC, despite the clear mitigation of ROS levels. This suggests that the increase of ROS levels is not a cause of the defective S-phase progression in *orc5-1 set1Δ*. Similarly, it has been shown that the DNA damage observed in *orc2-1* cells at high temperature is produced independently of ROS [51]. Therefore, we propose that the DNA damage is responsible for elevated ROS production [53] in the *orc5-1 set1Δ* mutant, and this increase in ROS in its turn accounts, at least in part, for the double mutant lethality.

## Discussion

Although H3K4 methylation may be required for full replication origins function [22], we were unable to detect any additive effect of Set1 loss on the minichromosome maintenance defect associated with *orc5-1* (Fig 2). This, and the fact that no global origin firing deficiency has been observed in *set1Δ* [31], strongly suggests that a synthetic reduction in origin activity is not sufficient to explain the *orc5-1 set1Δ* genetic interaction. Here, we presented evidence that this genetic interaction rather stems from a role for Set1 in replication fork progression that is unveiled when ORC function is affected. This conclusion is primarily based on the marked S phase lengthening manifested in *orc5-1 set1Δ* (Fig 3) and DNA damage accumulation (Fig 5 and S5 Fig).

The question arises about how exactly the *orc5-1* mutation renders the S phase sensitive to Set1 loss. Several evidences point to the intimate connection between the speed of replication forks and the frequency of origin activation. Altering replication fork speed trigger secondary responses in origins [54,55], and, conversely, decreasing the number of active origins induce compensatory increase in fork speed, partly due to higher dNTP availability [37,56]. Such compensatory increase in the rate of fork progression in *orc5-1* could explain our inability to detect a defect in S-phase progression through DNA content analysis. The premise that a faster replication fork progression is involved in the *orc5-1 set1Δ* genetic interaction is also supported by (i) the rescuing effect of lowering histone dosage that is known to limit fork speed [50], (ii) the negative effect of Sml1 loss on *orc5-1 set1Δ* viability (Fig 5C) given that an increase in the pool of dNTPs accelerates fork progression [56], and (iii) the recent identification of a negative control that H3K4 methylation exerts on fork velocity [32]. The absence of genetic interaction between *set1Δ* and *cdc17-1* (S1 Fig) can be considered as an additional argument as the *cdc17-1* mutation, which affects the lagging strand elongation, is expected to slow down the replication fork. In this setting, the negative control that Rad53 exerts on the rate of fork progression under replication stress conditions [57] could explain the sensitivity of *orc5-1* to Rad53 inactivation (Fig 4A and S4A Fig).

An observation that merits explanation is the negative impact of HU on the double mutant viability. This might seem unexpected since the primary effect of HU, the decrease of the dNTP pool (through RNR inhibition), could have improved the viability similarly to that of histone depletion. However, replication forks arrested by short-term HU treatment accumulate ssDNA [58], notably through the resection of nascent DNA [31]. In contrast, ssDNA formation is not detected at replication sites in cells with inhibited biosynthesis of the histones [50]. This difference in ssDNA accumulation could explain why, despite both reducing replication fork velocity, reducing the dNTP pool and depleting histone have opposite effect on *orc5-1 set1Δ* viability. Additionally, the fact that oxidative stress may contribute to the cytotoxic effect of HU [59] must be taken into account given the involvement of ROS in *orc5-1 set1Δ* lethality (Fig 9).

The local reduction of RPA-bound ssDNA at HU-stalled forks in *set1Δ* [31] seemingly contradicts the global increase of ssDNA observed in the *orc5-1 set1Δ*, estimated through the analysis of nuclear Rfa1 foci. Assuming that, as proposed [31], the loss of Set1 also effectively limits the nucleolytic degradation of nascent DNA in *orc5-1 set1Δ*, one simple explanation is that the restricted production of ssDNA at individual stalled fork is largely surpassed by an increase in the number of stalled forks. Alternatively, the forks whose progression is impaired in *orc5-1 set1Δ* may be not equivalent to HU-stalled forks and are not processed in the same way.

Meiotic DNA replication appears to be more sensitive to the combination of *orc5-1* and *set1Δ* mutations than mitotic cells at a temperature fully permissive for *orc5-1* (Fig 3B). Some characteristics of the meiotic S phase can explain this increased sensitivity such as the fact that replication origins are on average less frequently activated [60], and that the amount of dNTPs is more limited [61]. Thus, meiotic S phase can be considered to occur in mild replication stress conditions that would be equivalent to the presence of limited amounts of HU in mitotic cells. This illustrates how the physiological context can influence the degree to which replication fork progression is sensitive to the lack of Set1-dependent H3K4 methylation.

Considering the results of our study in the context of published work, we propose the following model for the *orc5-1 set1Δ* genetic interaction (Fig 10). Two consequences of the *orc5-1* mutation can be considered (Fig 10, middle panel). On one hand, as described previously [62], the activation of a Mad2-dependent pathway, possibly the SAC, seems primary responsible for the G2/M arrest that restrict cell division when ORC is defective, whereas a reduction in the number of functional replication forks compromises the activation of the DNA damage checkpoint during S phase [63]. The checkpoint initiating signal can be a defect in kinetochore-spindle attachment and/or in sister chromatid cohesion [40,41]. On the other hand, the weaker level of origin firing caused by *orc5-1* results in an increase of the replication fork speed [37]. This increase can potentially induce some replication stress [64], in particular at highly transcribed regions because of TRCs. The slowing down of the replicon by Set1-dependent H3K4 methylation, as it passes through highly expressed ORFs, turns out to be important to protect the genome from TRCs [32]. Accordingly, in the absence of Set1 (Fig 10, bottom panel), unrestrained fork progression caused by the reduction in origin firing in orc5-1, could lead to frequent TRCs, resulting in R-loops formation that will stall replication forks and thus impede S-phase progression. Lowering of the histone levels (*hht2-hhf2Δ*), by counteracting the fork rate increase due to *orc5-1* mutation, could therefore limit the occurrence of TRCs in absence of Set1. In contrary, TRCs are promoted by the removal of RNase H activity (*rnh1-rnh201Δ*) which stabilizes co-transcriptional R-loops.

**Fig 10.**
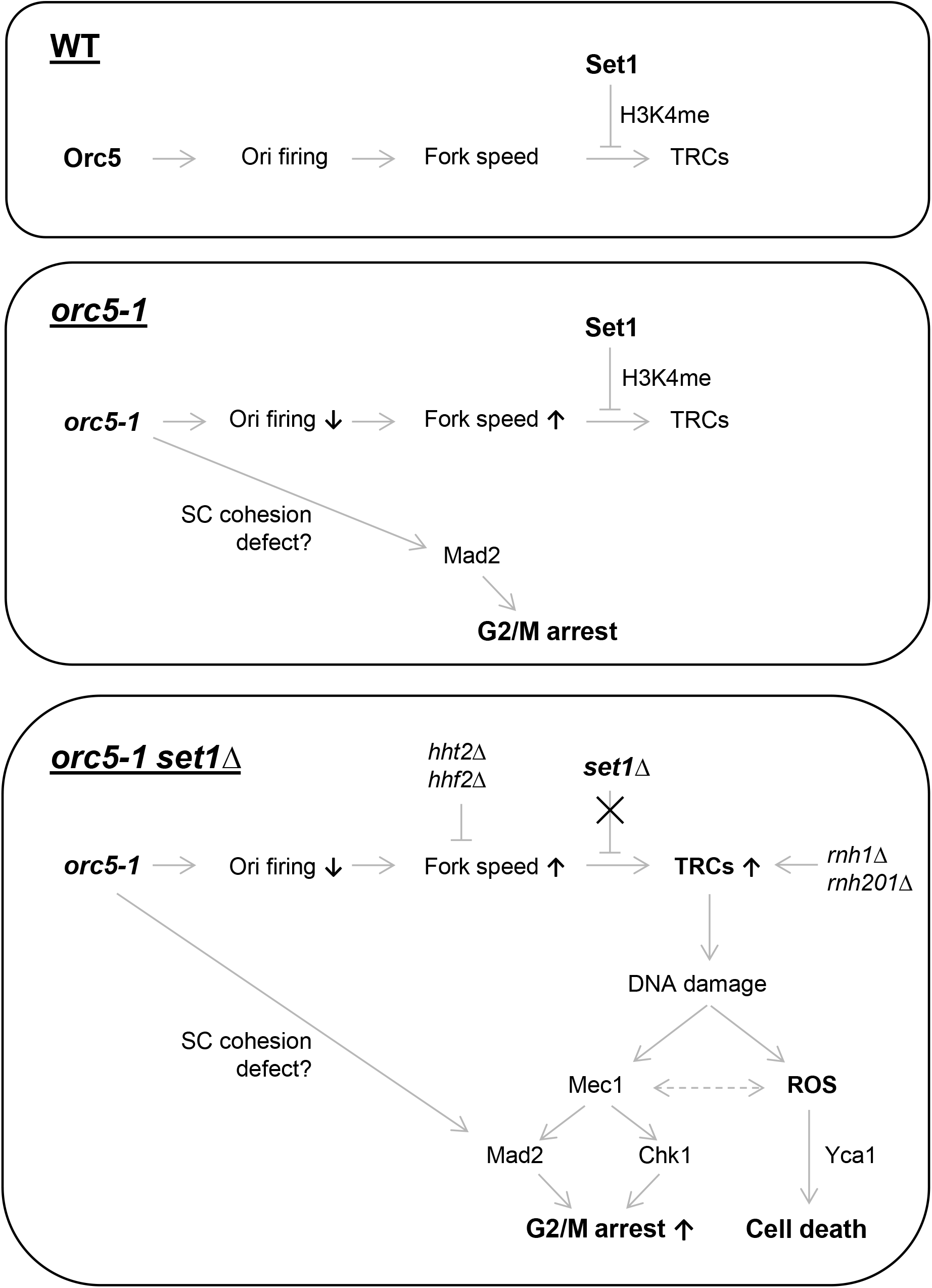
Proposed model for the genetic interaction between *orc5-1* and *set1Δ*. In the wild type (top), Orc5 ensures proper origin firing and Set1, through the transcription-deposited H3K4 methylation, helps to avoid the occurrence of TRCs at highly transcribed regions. The *orc5-1* mutation has two consequences (middle): first, the activation of Mad2 (via sister chromatid cohesion defect ?) that lead to a G2/M arrest, and, second, an increase of replication fork speed that compensates for weaker levels of origin firing. The increase in fork velocity is efficiently dampened by Set1-dependent H3K4 methylation, relieving highly transcribed regions from TCRs. In absence of Set1 (bottom), unrestrained fork progression at highly transcribed regions elevates the frequency of TRCs which are even more frequent if co-transcriptional R-loops are stabilized when RNase H activity is missing (*rnh1Δ rnh201Δ*). Conversely, lowering the histone levels (*hht2Δ hhf2Δ*) counteracts the fork rate increase due to *orc5-1* mutation, and thus relieves TRCs in absence of Set1. The DNA damage associated to TRCs reinforces the G2/M arrest, through activation of the Mec1-Chk1 signaling, and can induce a Yca1-dependent cell death, as a consequence of ROS production. See discussion for details.

R-loops are responsible for stalled forks at TRCs sites, and both are associated with ssDNA bound by RPA [4,10]. Nucleolytic processing of the ssDNA can generate DSBs, the resection of which can generate additional ssDNA required for their Rad52-dependent repair. Whatever its origin, the RPA-covered ssDNA is responsible for the excess of Rfa1 foci in *orc5-1 set1Δ* and activates Mec1 which, through both Mad2 and Chk1, reinforces the G2/M arrest of the *orc5-1* mutant (Fig 10, bottom panel). DNA damages, in the form of ssDNA and DSBs, are also a source of ROS [53]. Our data indicates that, above a certain threshold, ROS can induce a Yca1-dependent cell death. These two consequences of DNA damage - cell cycle arrest and ROS production - may be not independent as a mutual feed-forward relationship exists between Mec1 and ROS, with ROS production being partly dependent on Mec1 [51] and Mec1 activity requiring some ROS [65].

A role for Set1 in TRCs prevention was proposed in the specific context of checkpoint-defective (*rad53* mutants) cells treated with HU [32]. In this context, the fact that H3K4 methylation favors forks stalling at highly transcribed regions compromises their integrity, and the relief of this impediment to fork progression by ablating Set1 improves cell viability. Such a positive outcome contrasts with the negative impact of Set1 loss on cell viability when associated with the *orc5-1* mutation. This shows that the effect of Set1 ablation is context sensitive and can have opposite outcomes according to the way replication stress is induced. In checkpoint defective cells (*rad53* mutants) during an HU-induced stress, the structure and functionality of stalled forks is not preserved and relieving the impediment to fork progression due to H3K4 methylation is overall beneficial [32]. In checkpoint proficient cells (our study), the increase of fork velocity due to *orc5-1* favors the occurrence of TRCs and removing the protection provided by H3K4 methylation becomes detrimental. Our finding thus strengthens the notion that one major role of H3K4 methylation is to preserve the integrity of replication forks by modulating their velocity at the level of highly transcribed regions.

## Materials and methods

### Yeast strains and growth conditions

All strains used in this study are in the W303 or SK1 background and are listed in S1 Table. Standard conditions were used to grow and maintain strains on YPD (yeast extract-peptone-dextrose). Construction of *de novo* gene deletion strains was performed by PCR-mediated recombination and all double-mutant construction was performed by mating. For genetic complementation assays in *orc5-1 set1Δ* (Fig1A, middle), wild-type Set1 and mutant Set1G951S proteins fused to the Gal DNA binding domain were expressed under the control of the *ADH1* promoter from constructs integrated at the *trp1-1* locus as previously described [66].

To analyse meiotic S-phase, after growth in rich glucose medium (YPD), exponential phase cells were pregrown in rich acetate medium (YPA; 1% potassium acetate, 2% bacto-peptone, 1% bacto yeast extract supplemented with 25% amino acids) during 8 h, then diluted at 2 × 10^6^ cells/ml and grown in YPA during 14h. Cells were washed once with water and then inoculated into sporulation medium (1% potassium acetate supplemented with 25% amino acids) and incubated at 30° with vigorous agitation. For measure of sporulation levels, cells were directly striked from YPD plates on sporulation medium plates (1% potassium acetate supplemented with 25% amino acids). Sporulation rate was determined by counting asci visualized by light microscopy.

### Spore colony growth analysis

Tetrad dissection was performed on YPD plates using the MSM 400 dissection microscope (Singer Instrument Company) and isolated spores were incubated three days at 25°or two days at 30°. JPEG files of dissection plates were analyzed with ImageJ [67] to measure the area of each spore-derived colony.

### Temperature and DNA damage sensitivity assays

Freshly grown cells were taken, resuspended in water to 0.5 at DO_600_, and ten-fold serially diluted. Seven microliters of yeast cells at different dilutions were then spotted on YPD media or on YPD media containing either various concentrations of HU, NaCl (1M) or N-acetylcysteine (30mM). Images were taken after incubation at the indicated temperatures for 2-5 days.

### Plasmid maintenance assays

Plasmid maintenance assays were performed as described previously [22]. Yeast strains containing a *CEN4/ARS1/URA3* plasmid (pRM102, [68] were grown to log phase in selective medium (lacking uracil) and 100–200 cells were plated on both selective and nonselective media to establish an initial percentage of plasmid-bearing cells. These cultures were also diluted to a concentration of 1 × 10^5^ cells/ml in 5 ml of nonselective media and grown for 8–10 generations before once again plating on both selective and nonselective media. Precise generation numbers were calculated using the following formula: *n* = log(*C*_F_/*C*_I_)/log(2), where *C*_F_ represents the final number of cells as measured by OD_600_ and *C*_I_ represents the starting cell number of 10^5^ cells/ml. After 2 days of growth, colonies were counted, and the plasmid loss rate (*L*) per generation (*n*) was calculated using the following formula: *L* = 1 − (%*F*/%*I*)^(1/*n*)^, where %*F* is the final percentage of cells that retained the plasmid and %*I* is the initial percentage of cells that contain the plasmid.

### Analysis of DNA content

Cells were fixed in 70% ethanol. After rehydration in PBS, the samples were incubated at least 2 h with RNase A (1 mg/ml) at 37°. Cells were resuspended in 50 μg/ml propidium iodide in PBS for at least 15 min at room temperature. After a wash in PBS, cells were resuspended in 5 μg/ml propidium iodide, sonicated briefly to remove cell clumps, and the DNA content was determined by FACS with a Becton Dickinson FACSCalibur.

### Cell-Cycle Progression and BrdU Labeling

Yeast cultures were grown at 25°to an A_600_ of 0.6 and arrested for 2-3 hours in G_1_ with α factor (5 μg/ml), 1-2 hours at 25° then 1 hour at 37°. Cells were washed two times with distilled water and one time with fresh YPD medium and released into the first cell cycle in YPD. To label newly replicated DNA, BrdU (Sigma) was added to the medium to a final concentration of 100 μg/ml just after the release from the G1 block. Two cell samples were collected for each time point. One sample (1ml) was used to monitor cell-cycle progression by flow cytometry (see Analysis of DNA content). The other sample (14ml) was used to extract genomic DNA. The cell pellet was resuspended in 200 μl TES (Tris EDTA + 1% SDS) together with 200 μl of glass beads and 400 μL of PCI (Phenol:Chloroform:Isoamyl alcohol) before vigorous vortexing for 30 min in a multi vortex. After spinning at 13.000 rpm for 5 min, 1ml cold (−20°) absolute ethanol was added to the aqueous layer. After rapid mixing, microtubes were put on ice 5 min before spinning at 13.000 rpm for 10 min at 4°. Pellets were washed with ethanol 70% before spinning again. Dry pellets were resuspended in 50 to 100 μl of TE-RNase (Tris EDTA + RNAse A). After 30-40 min at 50°, genomic DNA (2-3 μl) was checked on 0.8% agarose gel.

### Detection and Quantification of BrdU

Genomic DNA (20 μl, see previous section) was denaturated in 1.5 N NaOH, 3M NaCl for 10 min at room temperature and spotted directly onto a nitrocellulose membrane (Protran). The membrane was crosslinked with UV (1200J/m^2^) and neutralized with 0.5 M Tris/HCl (pH 7.5), 0.5 M NaCl (20 min at room temperature). The membrane was blocked for 20 min at room temperature in TBS 1X, 0.1% Tween-20, 5% milk. The membrane was then incubated with a monoclonal antibody against BrdU (1:4000; Abcam ab12219) overnight at 4° in TBS1X, 0.1% Tween-20, 0.1% milk. An Alexa Fluor 680 anti-mouse goat IgG (1:5000; Molecular Probes) was used as a secondary antibody. Emitted fluorescence of the secondary antibodies was detected by using an Odyssey Imager (LI-COR Bioscience). The signal of the dots was quantified using the program Image J1.32.

### Fluorescence microscopy

Cells expressing Rfa1 tagged with cyan fluorescent protein (CFP) or Rad52 tagged with yellow fluorescent protein (YFP) where grown in liquid YPD (+ adenine) medium to exponential phase at 30°, harvested, washed in PBS, and placed on a glass side. Observations of cells were performed using a Nikon Eclipse Ti microscope with a 100x oil immersion objective. Cell images were captured with a Neo sCMOS Camera (Andor). Fluorophores were visualized using band-pass CFP (Rfa1-CFP) or YFP (YFP-Rad52) filter sets. For each field of view, a single DIC image and 11 CFP or YFP images at 0.3μM intervals along the Z-axis were acquired. CFP and YFP foci were visualized and quantified using ImageJ software. For each strain, at least 200 cells were scored for RFa1-CFP or Rad52-YFP foci. Mean fluorescence intensity within constant square regions placed in the nucleus outside the foci area was measured to get the average nuclear background fluorescence (N). Quantification of cell morphology was determined by analysis of at least 200 cells at each time point.

### Measurement of ROS levels

After an overnight incubation at 25°, cells were inoculated (A_600_ of 0.2) in fresh YPD medium (supplemented with adenine) and grown at 25° or 32° during seven hours, with or without NAC (30mM). About 10^7^ cells were collected, centrifugated, resuspended in 100μl Tris EDTA 50mM pH7.5 with 0.1μl CellROX^®^ Green reagent (Life Technologies) and incubated during 30 min at the same temperature of 25° or 32°. Intracellular levels of reactive oxygen species (ROS) was assessed by measuring the mean fluorescence using flow cytometry with a Becton Dickinson FACSCalibur. Control autofluorescence signals were obtained by incubating cells in absence of CellROX^®^ Green reagent.

### Data availability

All strains listed on S1 Table that were generated in the V.G. lab are available upon request.

## Acknowledgements

We thank Valérie Garcia, Christelle Cayrou and Dmitri Churikov for critical reading of the manuscript; Morgane Joessel and Gaëtan Vaccari for their contribution in this work during their training course in the laboratory; Michael Weinreich for the *orc5-1*, *cdc6-4*, *mcm2-1* and *cdc17-1* strains; Akash Gunjan for strains harboring the *hht1-hhf1Δ* and *hht2-hhf2Δ* loci; Nicanor Austriaco for the strain harboring the *yca1Δ* locus.

## Supporting information

**S1 Table.**
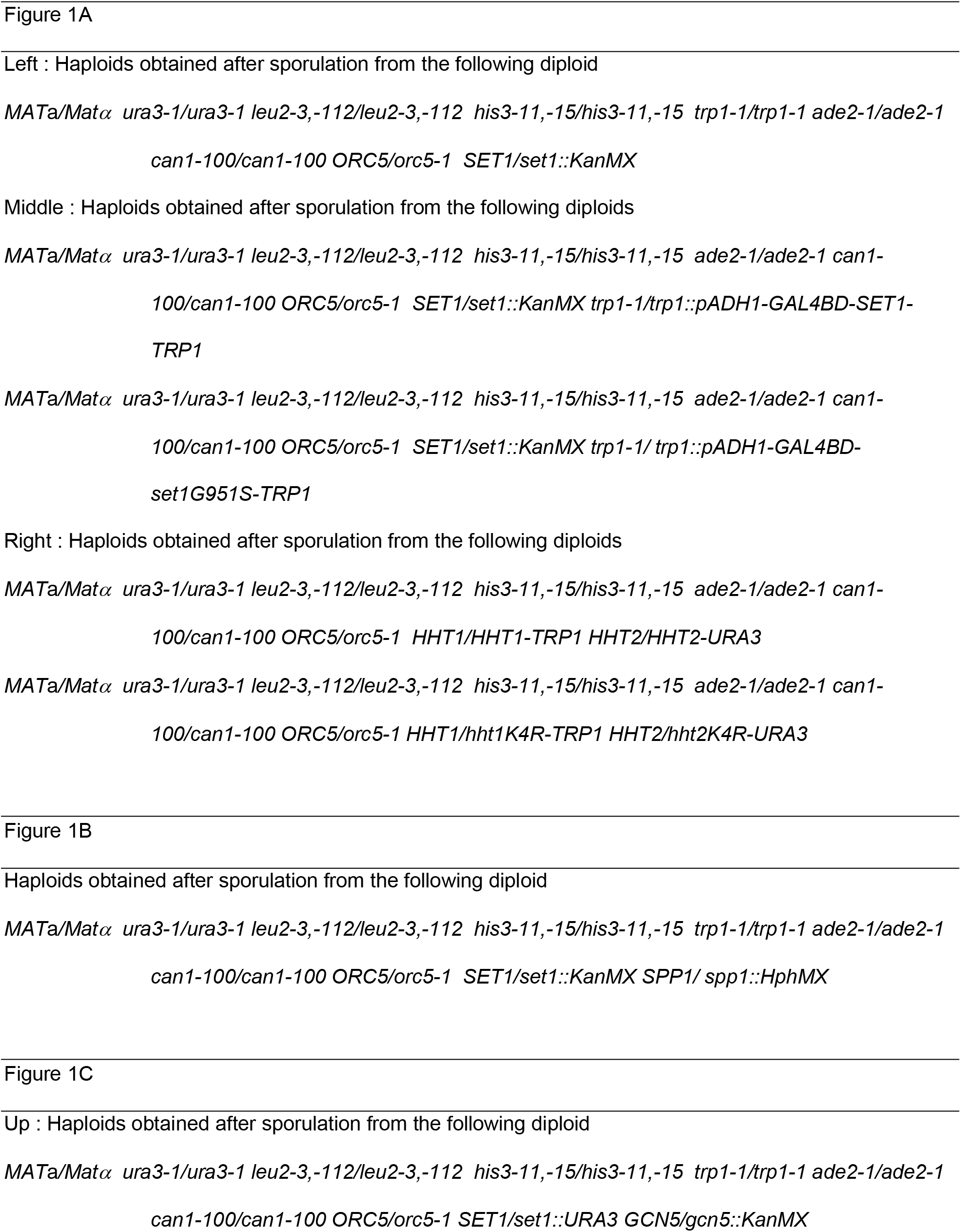
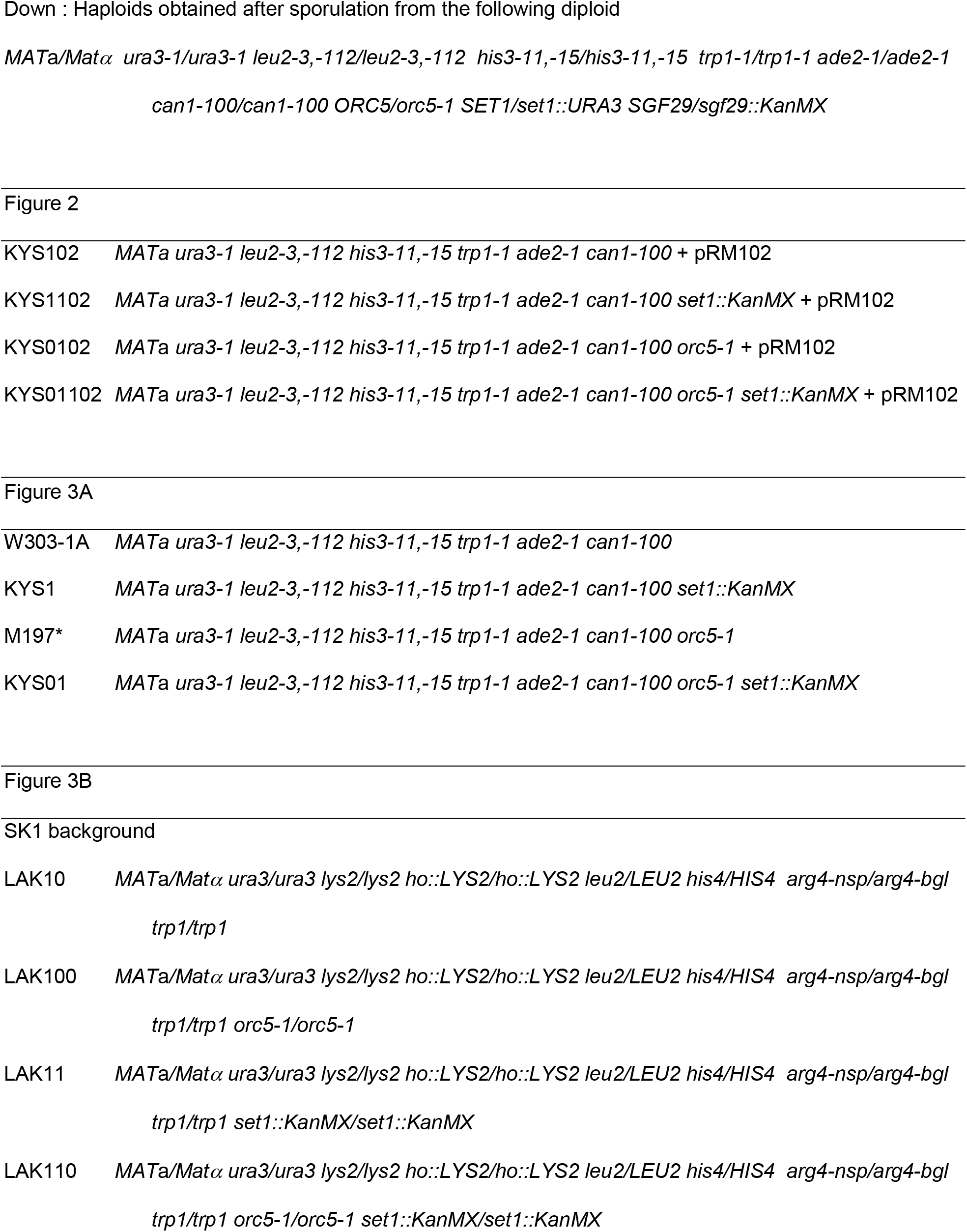
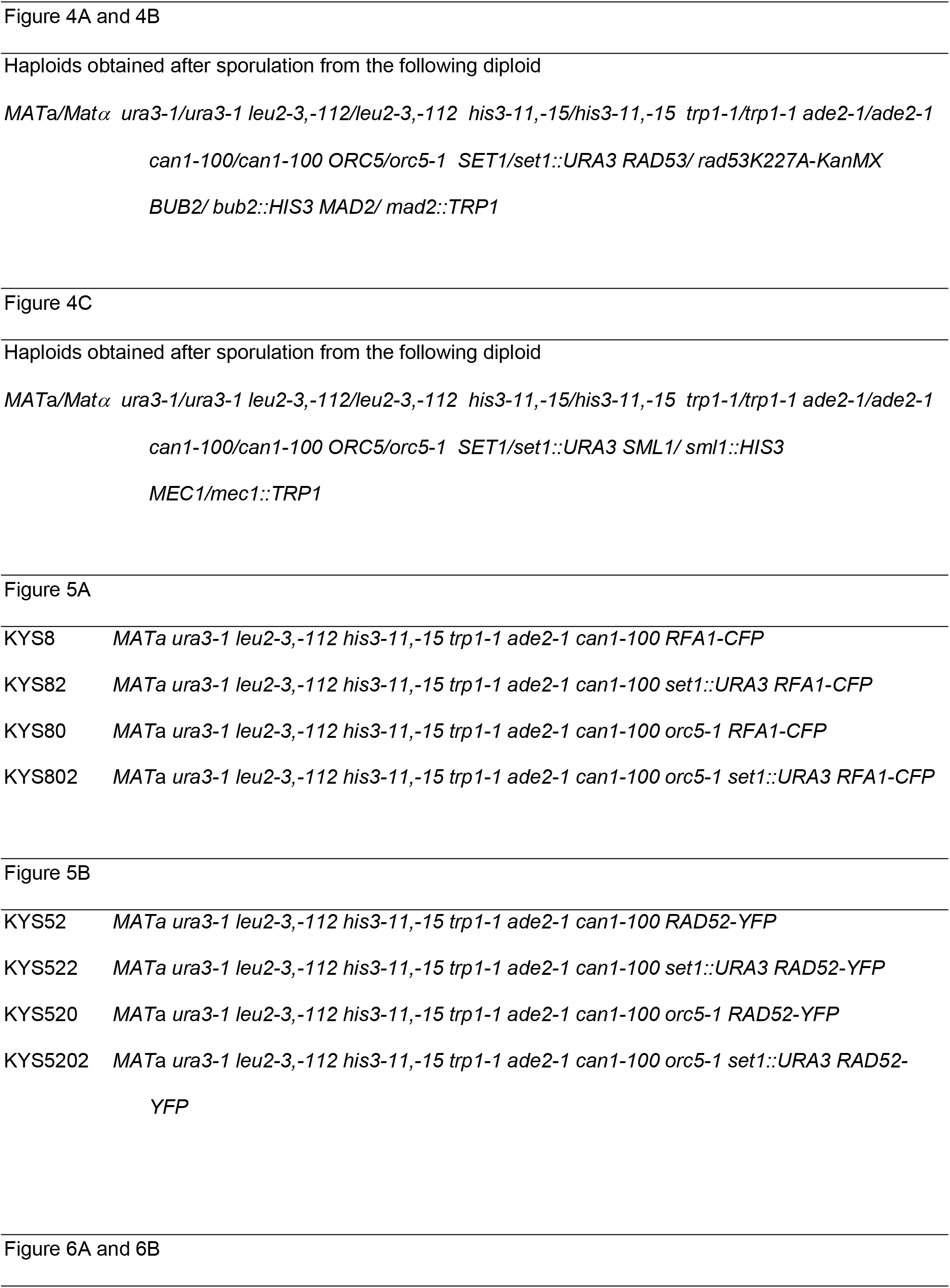
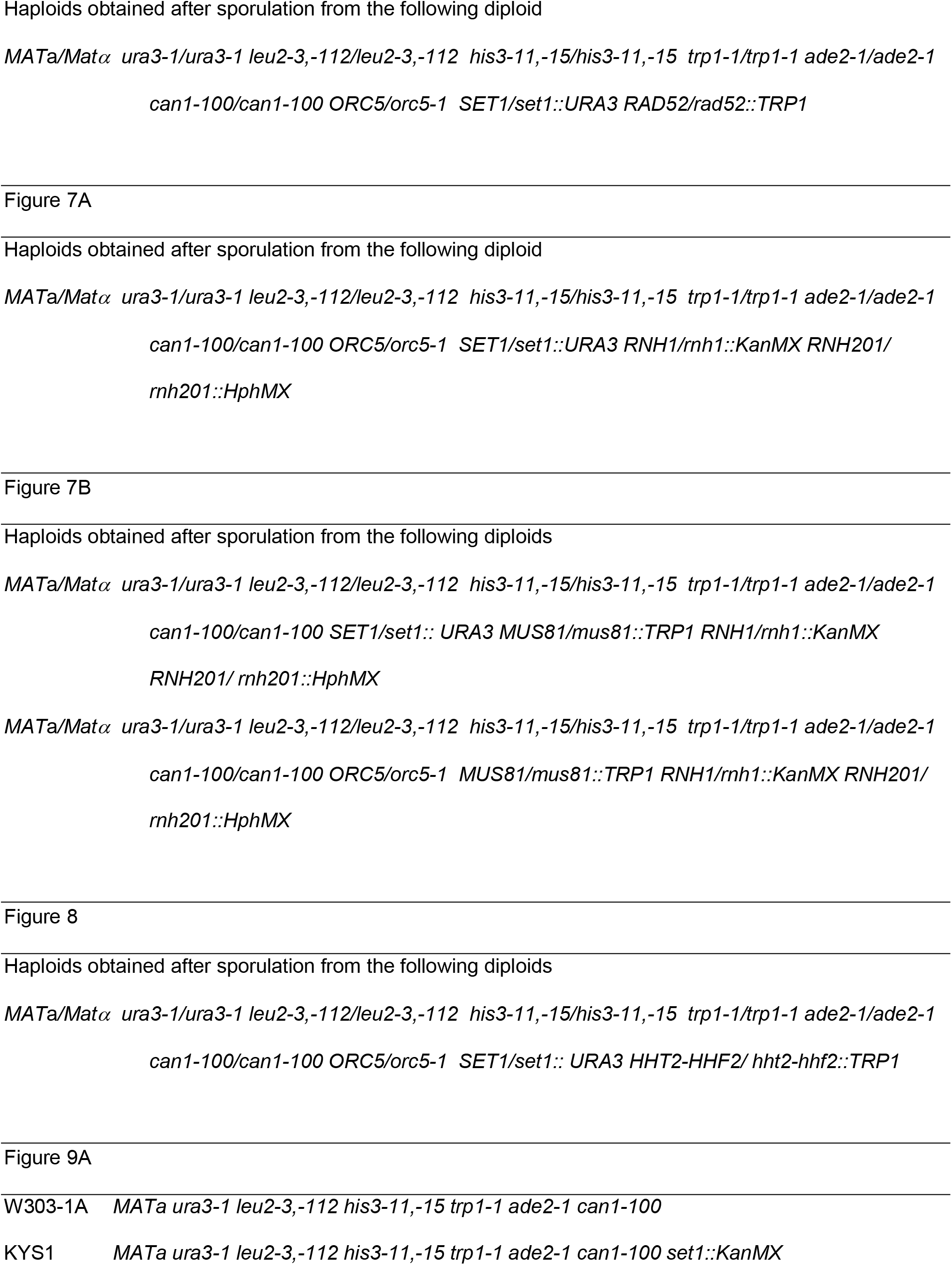
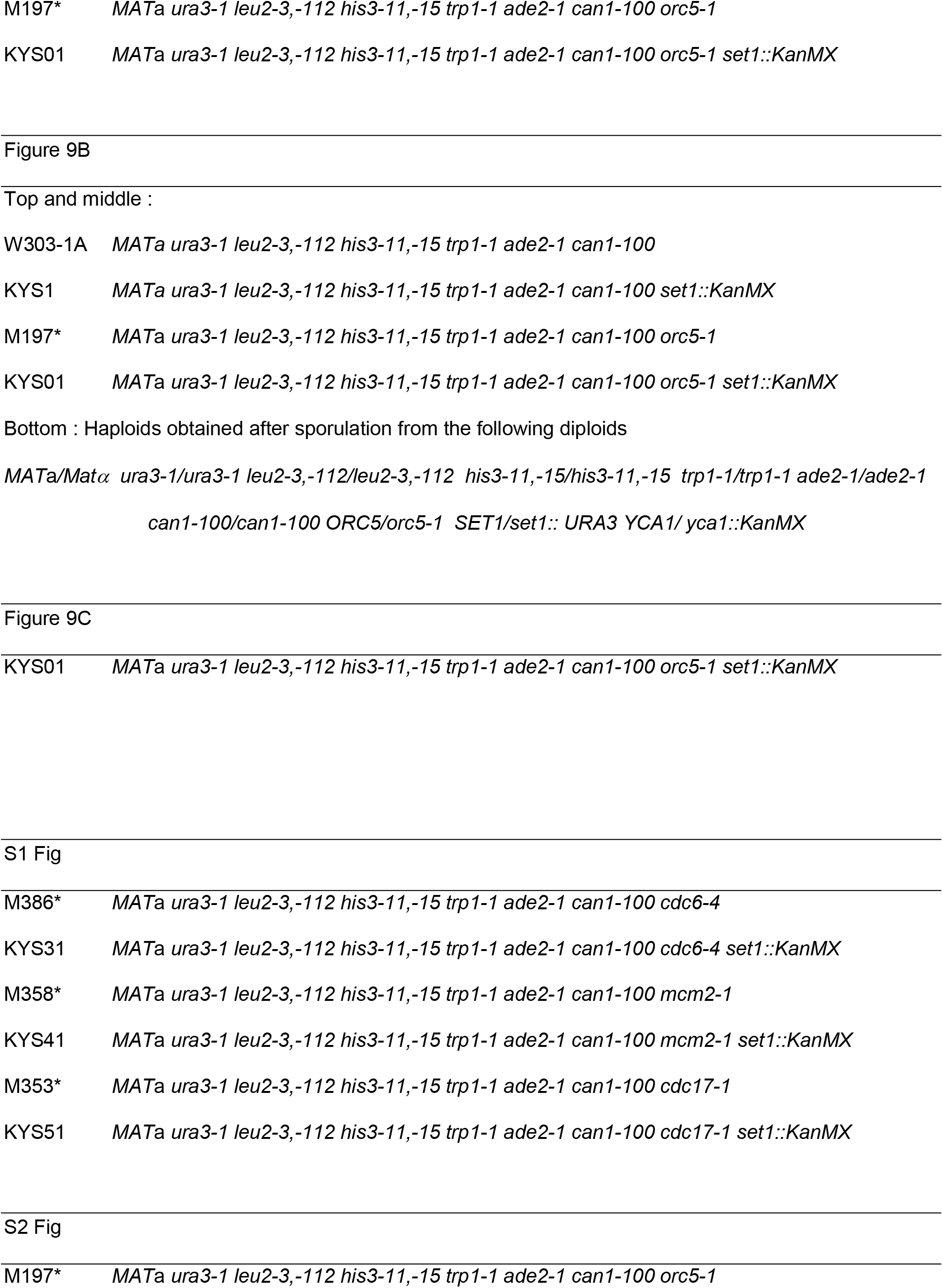
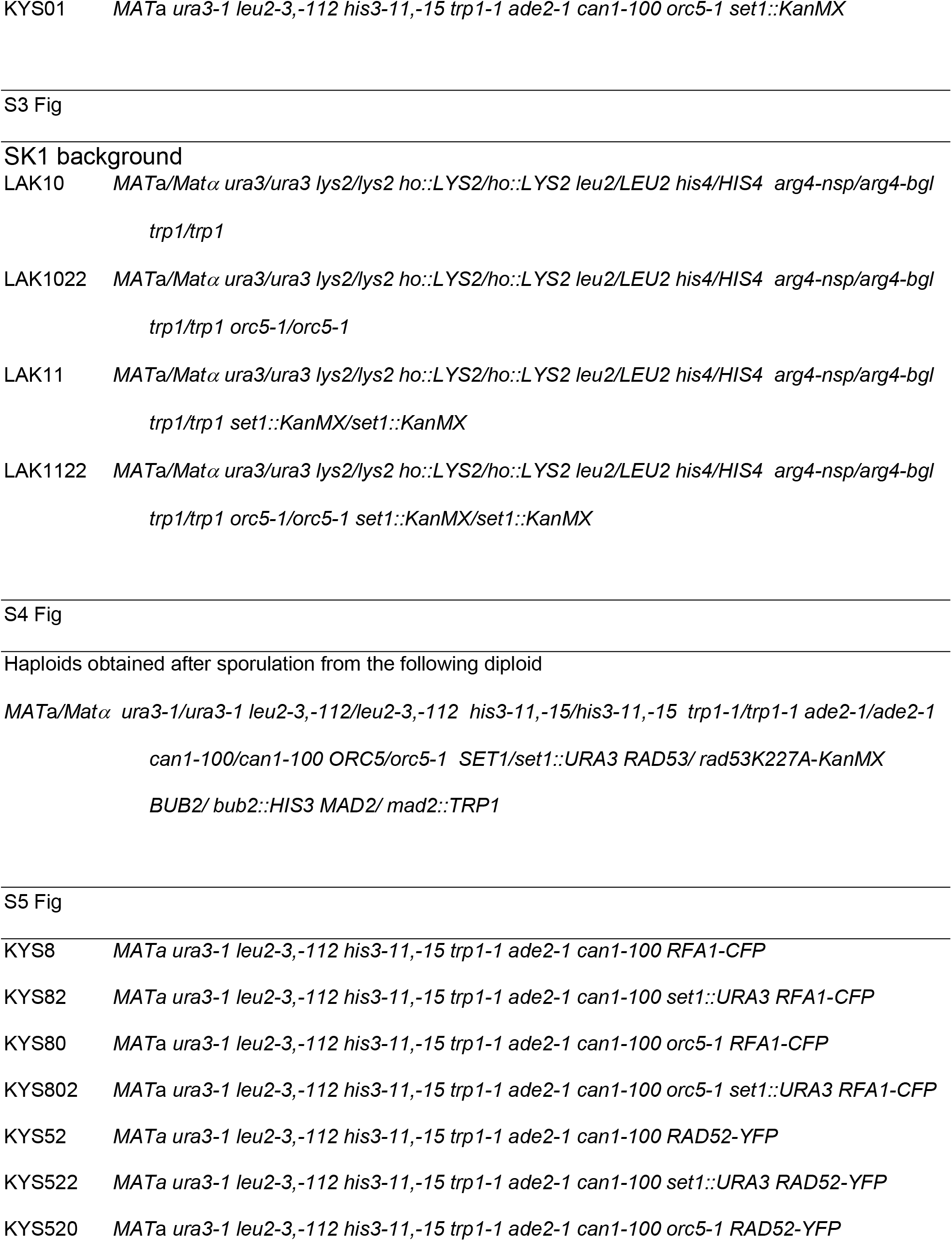
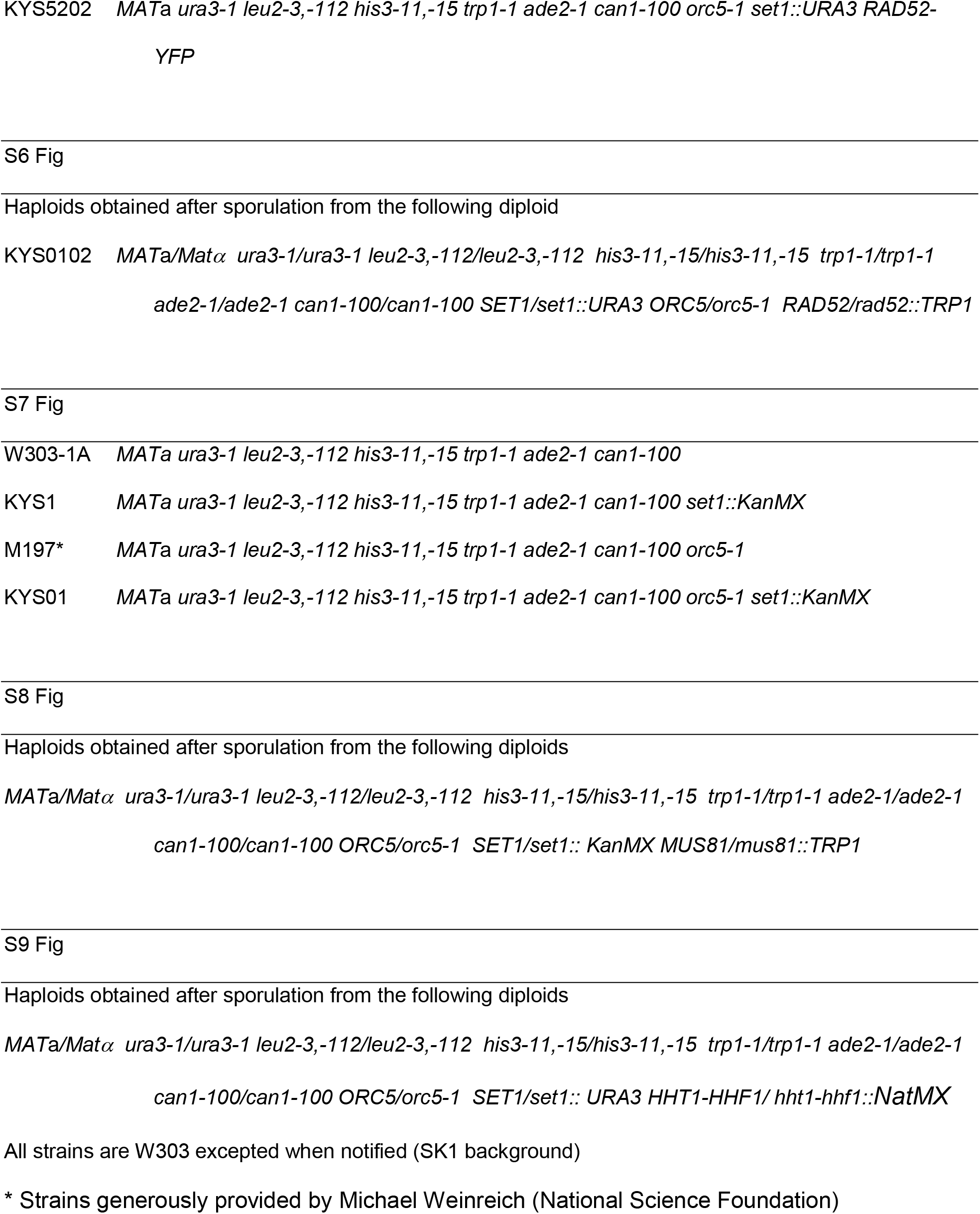
Strains used in this study.

**S1 Fig.**
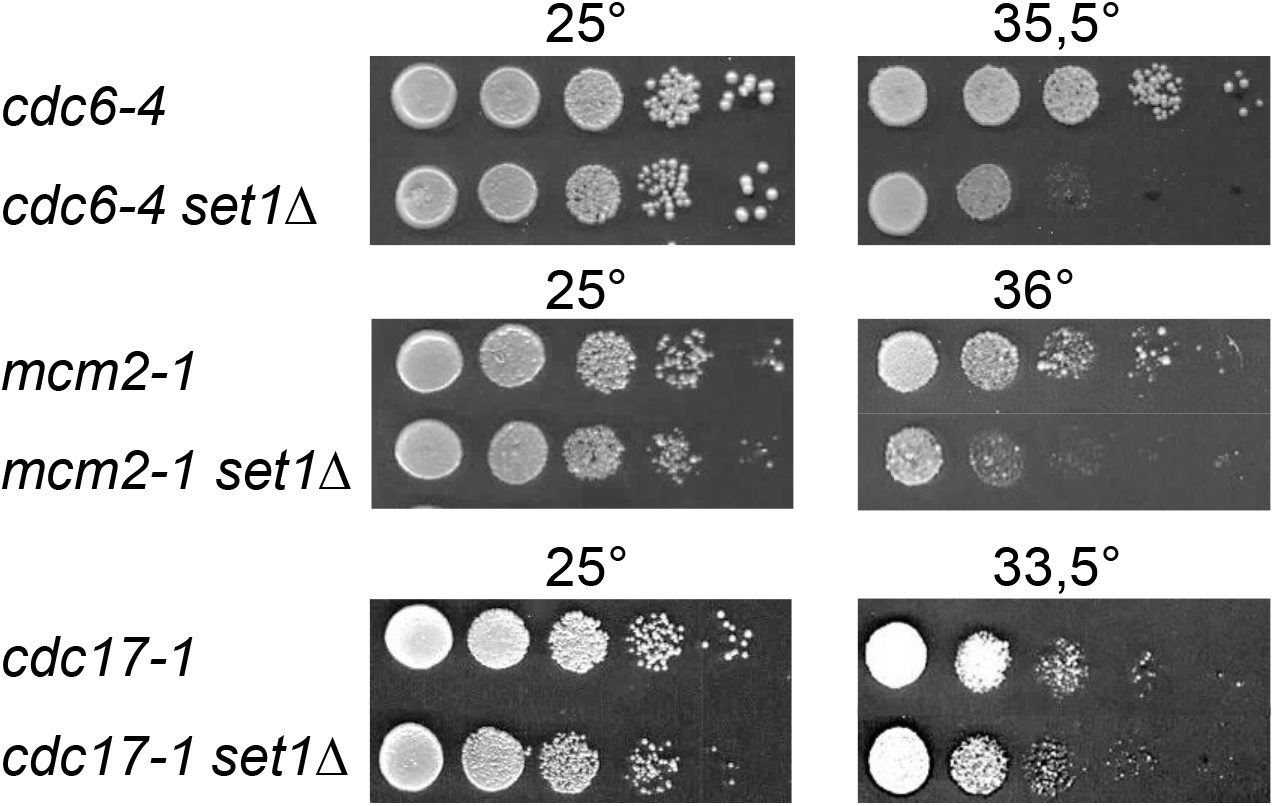
Impact of Set1 loss on the thermosensitivity of *cdc6-4*, *mcm2-1* and *cdc17-1*. Tenfold serial dilutions of the respective strains were spotted onto YPD and incubated at the indicated temperatures for 3-4 days.

**S2 Fig.**
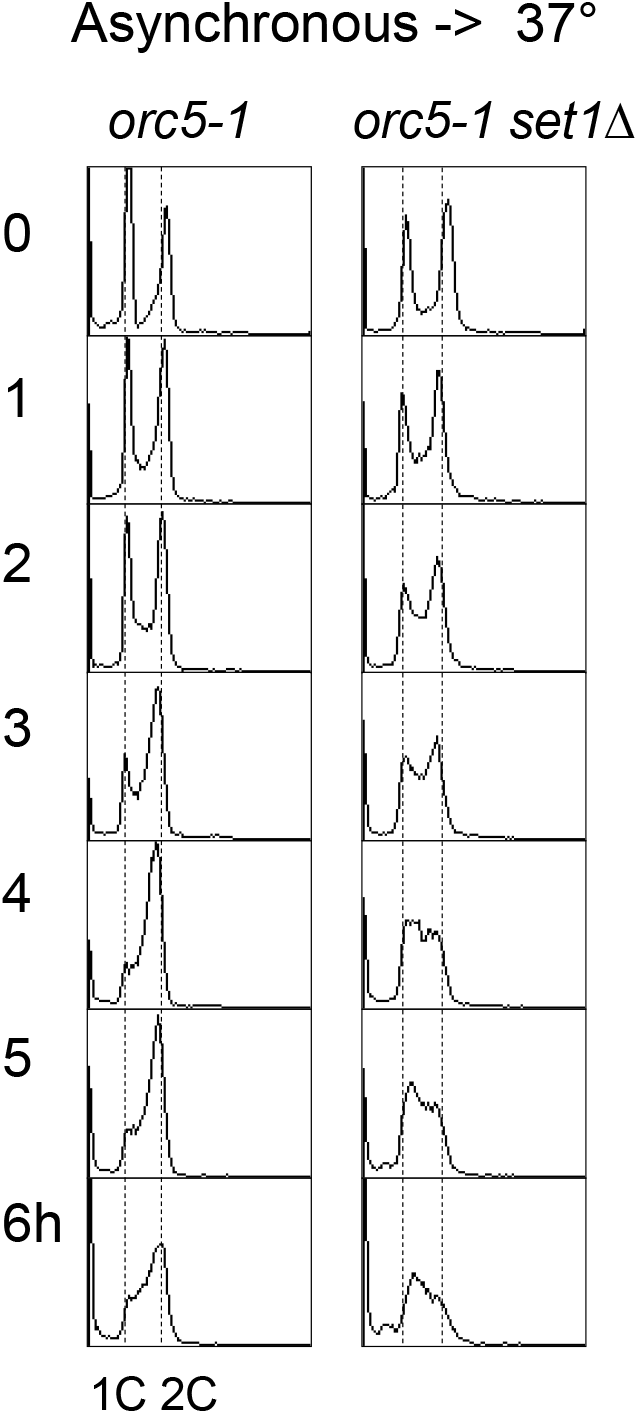
The S phase becomes limiting in *orc5-1 set1Δ* at 37°. Unsynchronized cultures of *orc5-1* and *orc5-1 set1Δ* growing at 25° were shifted to 37°. DNA content was determined at the indicated intervals (hours). 1C and 2C (broken lines) indicate the DNA content of cells with unreplicated or fully replicated DNA, respectively.

**S3 Fig.**
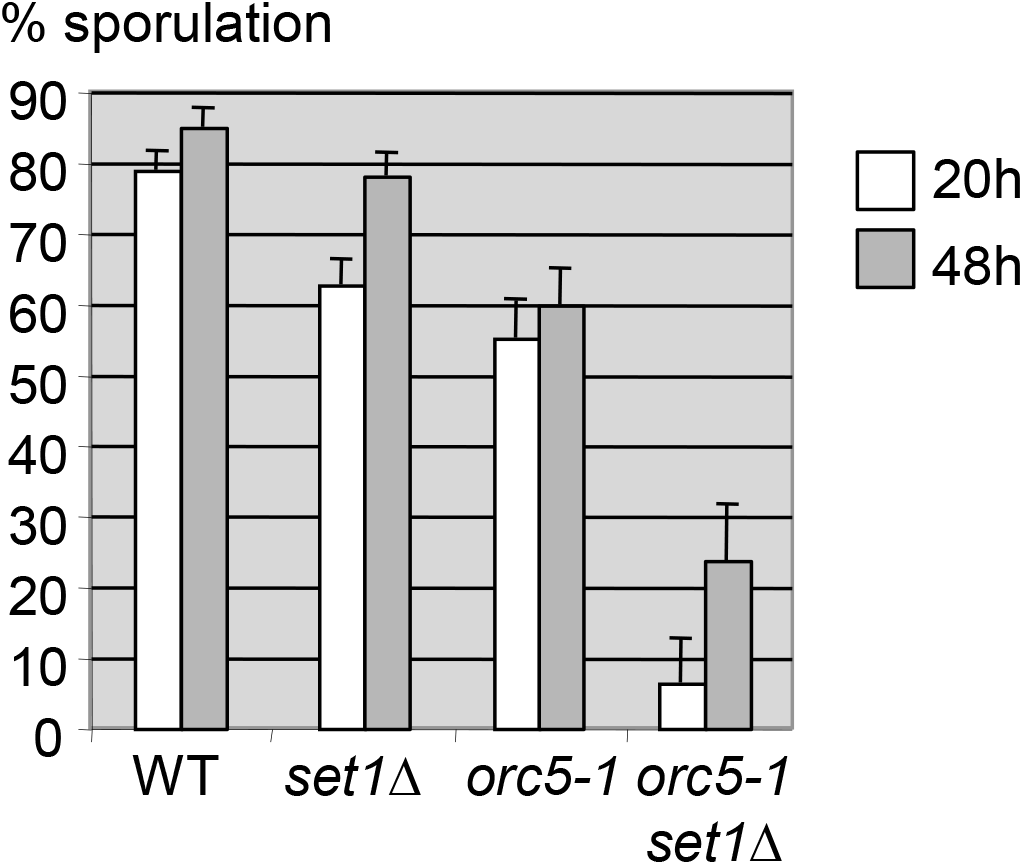
Low sporulation levels in the *orc5-1 set1Δ* SK1 diploid. Percentage of sporulation of the indicated diploids after 20h or 48h on sporulation plates at 25°.

**S4 Fig.**
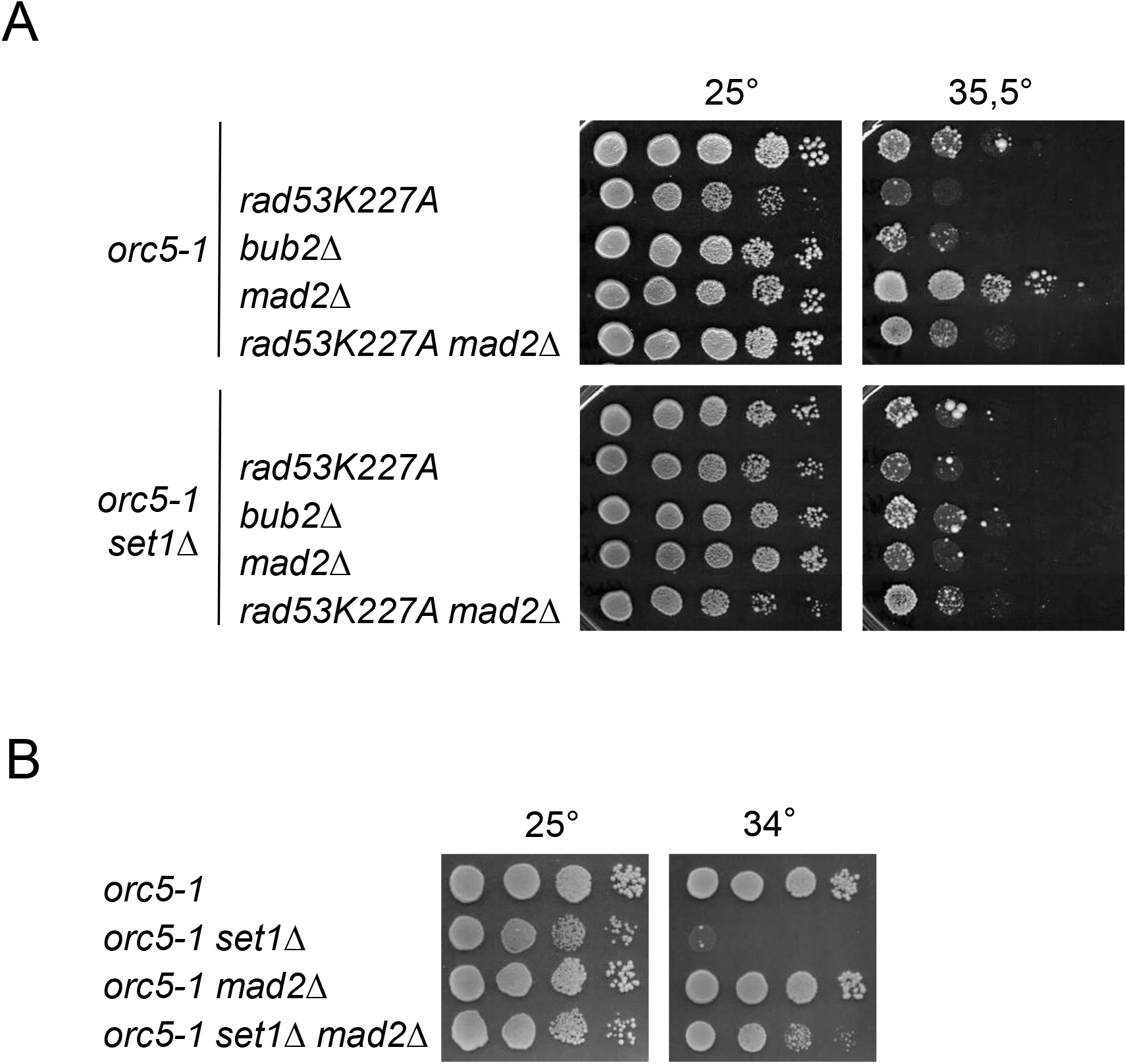
Differential impact of *mad2Δ* on *orc5-1* and *orc5-1 set1Δ* thermosensitivity. Tenfold serial dilutions of the respective strains were spotted onto YPD plates and incubated at the indicated temperatures for 3-4 days. (A) Same experiment as in Figure 4A with independent clones. (B) A rescue effect of *mad2Δ* on *orc5-1 set1Δ* viability is seen at the lower temperature of 34°.

**S5 Fig.**
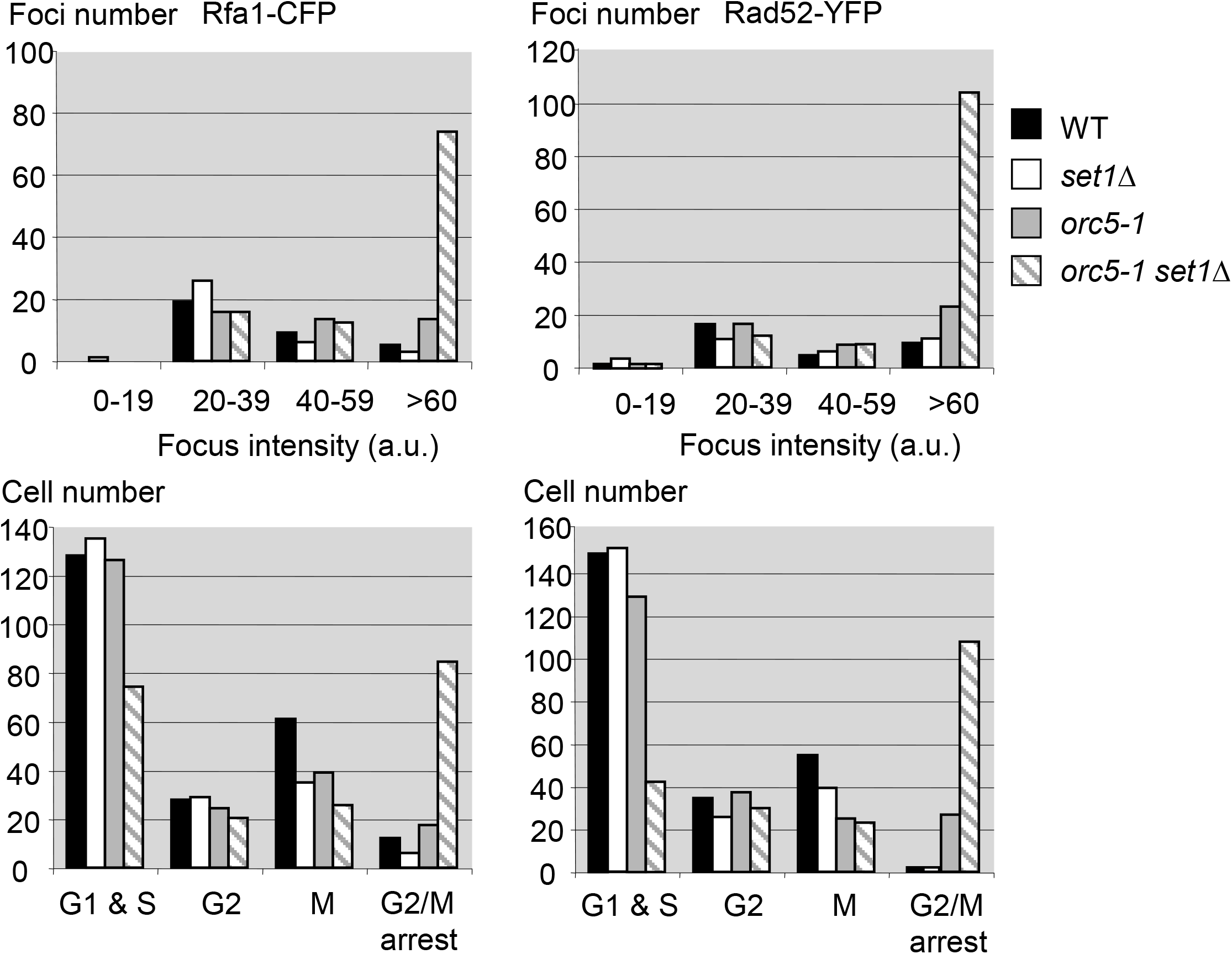
Heightened spontaneous DNA damage levels in *orc5-1 set1Δ* cells. Top, ranking of spontaneous Rfa1-CFP (left) or Rad52-YFP (right) foci according to their intensity in exponentially growing cells (30°) of the indicated genotypes (a.u.: arbitrary units). Bottom, distribution of the cells according to the cell cycle stage.

**S6 Fig.**
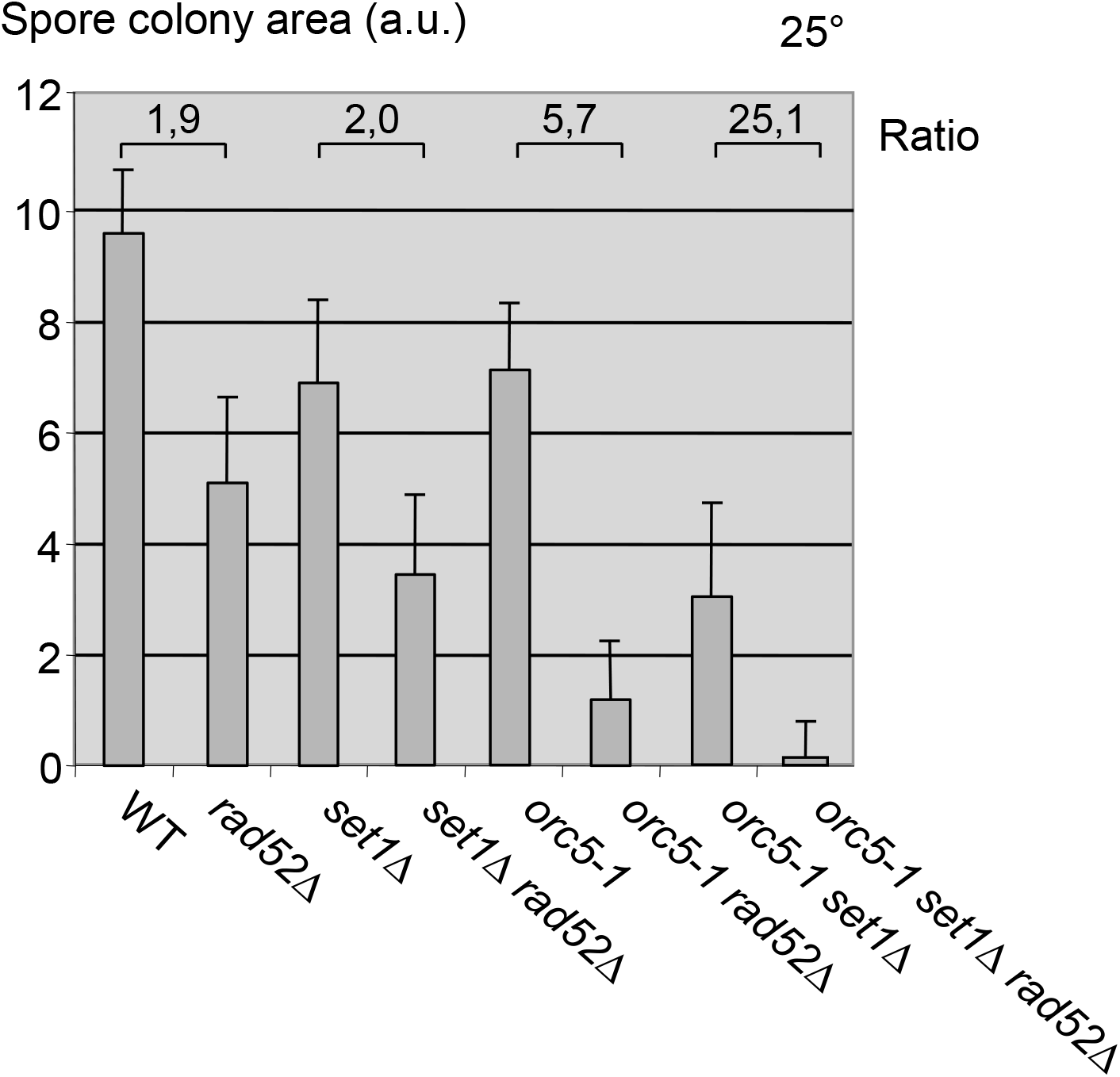
Growth of *orc5-1 set1Δ* colonies is greatly dependent on Rad52. The area of single-spore derived colonies was determined after three days at 25° since spore isolation (a.u.: arbitrary units). The ratio of the mean area of *RAD52* and *rad52Δ* colonies for each genotype is indicated.

**S7 Fig.**
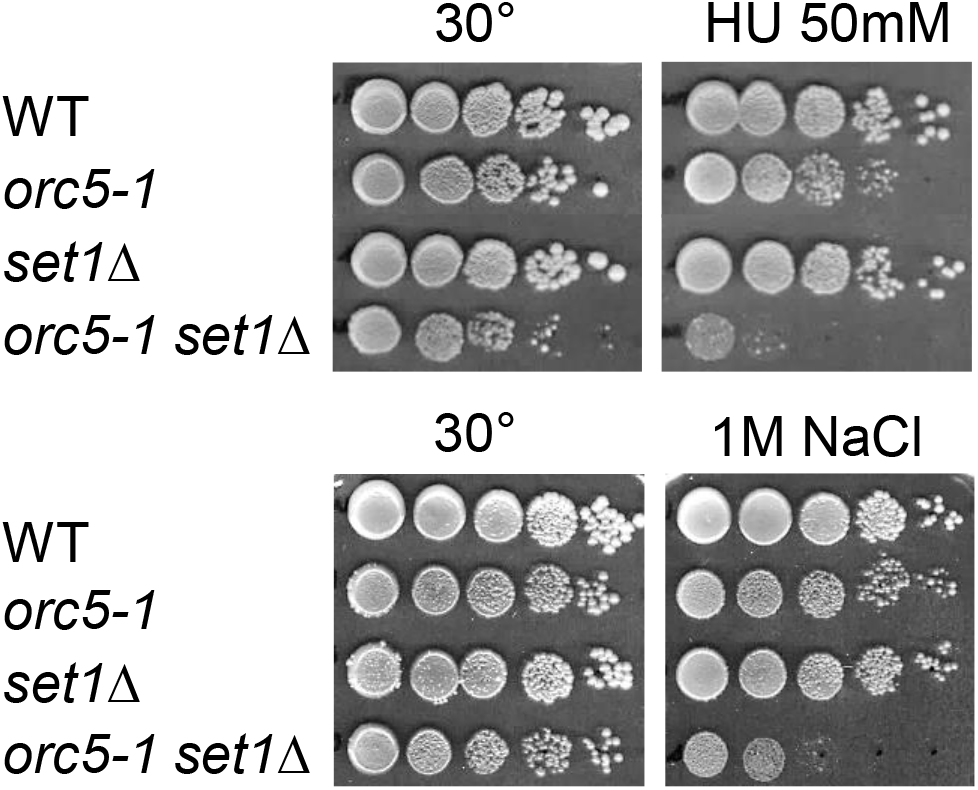
Hypersensitivity of *orc5-1 set1Δ* to HU and osmostress. Tenfold serial dilutions of the respective strains were spotted onto YPD plates containing HU or NaCl at the indicated concentrations and incubated at 30° for 4 days.

**S8 Fig.**
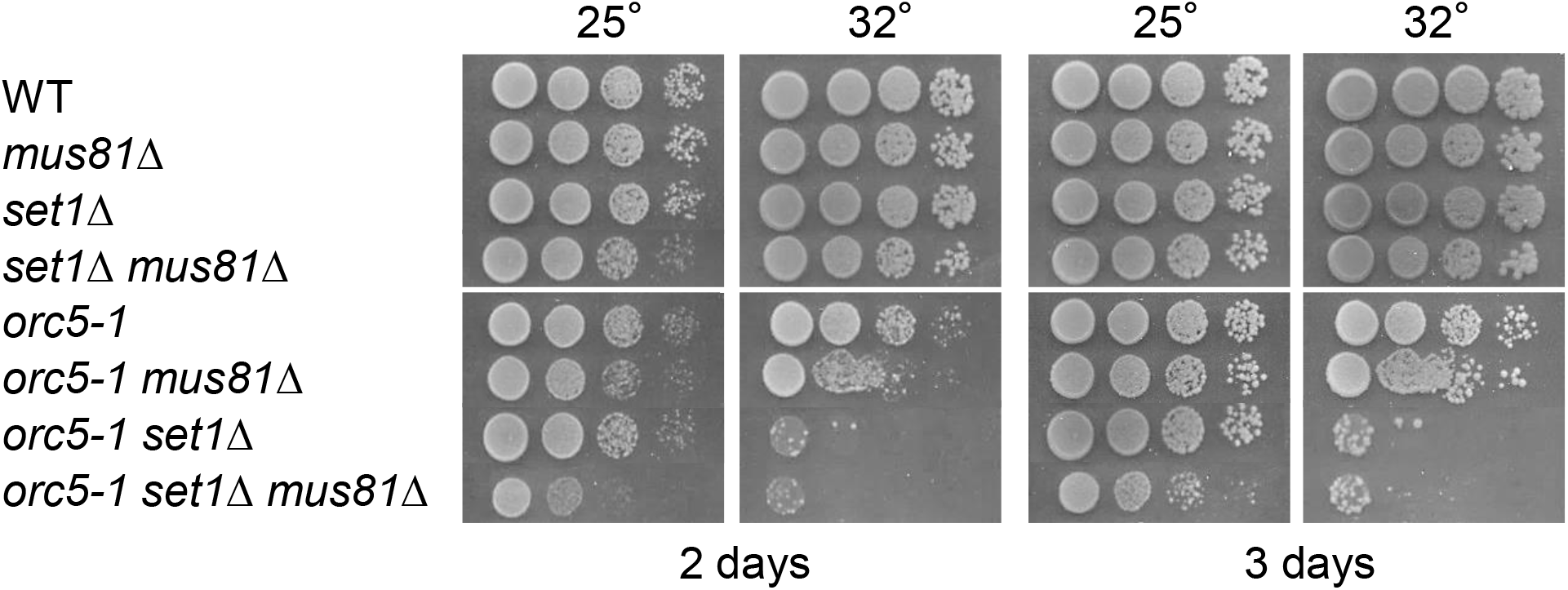
Limited effect of Mus81 loss. Tenfold serial dilutions of the respective strains were spotted onto YPD plates and incubated at the indicated temperatures for 2 and 3 days.

**S9 Fig.**
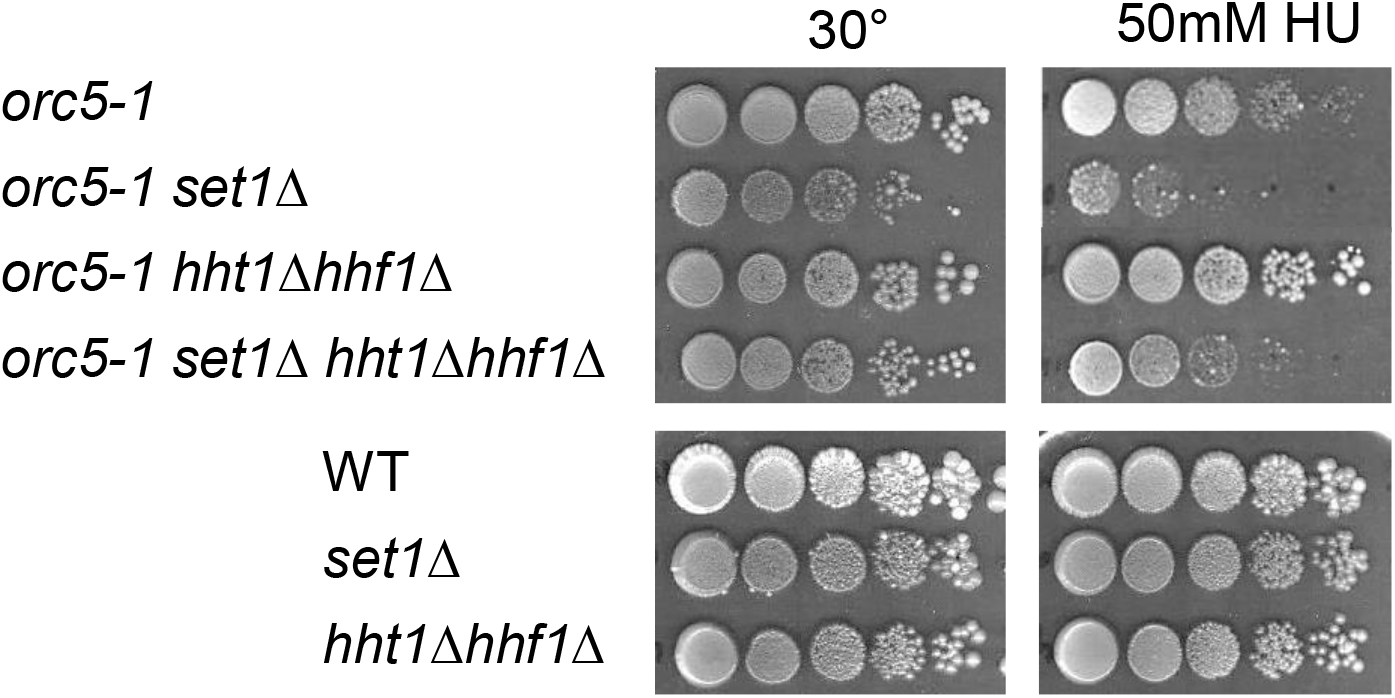
Effect of reducing histone gene dosage through *HHT1-HHF1* deletion. Tenfold serial dilutions of the respective strains were spotted onto YPD plates without or with HU (50mM) and incubated at 30° for 3-4 days.

## References

1. Bell SP, Labib K. Chromosome Duplication in Saccharomyces cerevisiae. Genetics. 2016;203: 1027–1067. doi:10.1534/genetics.115.186452

2. Gómez-González B, Aguilera A. Transcription-mediated replication hindrance: a major driver of genome instability. Genes Dev. 2019;33: 1008–1026. doi:10.1101/gad.324517.119

3. Hamperl S, Bocek MJ, Saldivar JC, Swigut T, Cimprich KA. Transcription-Replication Conflict Orientation Modulates R-Loop Levels and Activates Distinct DNA Damage Responses. Cell. 2017;170: 774–786.e19. doi:10.1016/j.cell.2017.07.043

4. Nguyen HD, Yadav T, Giri S, Saez B, Graubert TA, Zou L. Functions of Replication Protein A as a Sensor of R Loops and a Regulator of RNaseH1. Mol Cell. 2017;65: 832–847.e4. doi:10.1016/j.molcel.2017.01.029

5. Sabatinos SA, Forsburg SL. Managing Single-Stranded DNA during Replication Stress in Fission Yeast. Biomolecules. 2015;5: 2123–2139. doi:10.3390/biom5032123

6. Zou L, Elledge SJ. Sensing DNA damage through ATRIP recognition of RPA-ssDNA complexes. Science. 2003;300: 1542–1548. doi:10.1126/science.1083430

7. Shechter D, Costanzo V, Gautier J. Regulation of DNA replication by ATR: signaling in response to DNA intermediates. DNA Repair (Amst). 2004;3: 901–908. doi:10.1016/j.dnarep.2004.03.020

8. Weinert TA, Kiser GL, Hartwell LH. Mitotic checkpoint genes in budding yeast and the dependence of mitosis on DNA replication and repair. Genes Dev. 1994;8: 652–665. doi:10.1101/gad.8.6.652

9. Moriel-Carretero M, Aguilera A. A postincision-deficient TFIIH causes replication fork breakage and uncovers alternative Rad51- or Pol32-mediated restart mechanisms. Mol Cell. 2010;37: 690–701. doi:10.1016/j.molcel.2010.02.008

10. Allen C, Ashley AK, Hromas R, Nickoloff JA. More forks on the road to replication stress recovery. J Mol Cell Biol. 2011;3: 4–12. doi:10.1093/jmcb/mjq049

11. Eaton ML, Galani K, Kang S, Bell SP, MacAlpine DM. Conserved nucleosome positioning defines replication origins. Genes Dev. 2010;24: 748–753. doi:10.1101/gad.1913210

12. Hizume K, Yagura M, Araki H. Concerted interaction between origin recognition complex (ORC), nucleosomes and replication origin DNA ensures stable ORC-origin binding. Genes Cells. 2013;18: 764–779. doi:10.1111/gtc.12073

13. Kurat CF, Yeeles JTP, Patel H, Early A, Diffley JFX. Chromatin Controls DNA Replication Origin Selection, Lagging-Strand Synthesis, and Replication Fork Rates. Mol Cell. 2017;65: 117–130. doi:10.1016/j.molcel.2016.11.016

14. Ramachandran S, Henikoff S. Replicating Nucleosomes. Sci Adv. 2015;1. doi:10.1126/sciadv.1500587

15. Bellush JM, Whitehouse I. DNA replication through a chromatin environment. Philos Trans R Soc Lond, B, Biol Sci. 2017;372. doi:10.1098/rstb.2016.0287

16. Espinosa MC, Rehman MA, Chisamore-Robert P, Jeffery D, Yankulov K. GCN5 is a positive regulator of origins of DNA replication in Saccharomyces cerevisiae. PLoS ONE. 2010;5: e8964. doi:10.1371/journal.pone.0008964

17. Unnikrishnan A, Gafken PR, Tsukiyama T. Dynamic changes in histone acetylation regulate origins of DNA replication. Nat Struct Mol Biol. 2010;17: 430–437. doi:10.1038/nsmb.1780

18. Masumoto H, Hawke D, Kobayashi R, Verreault A. A role for cell-cycle-regulated histone H3 lysine 56 acetylation in the DNA damage response. Nature. 2005;436: 294–298. doi:10.1038/nature03714

19. Driscoll R, Hudson A, Jackson SP. Yeast Rtt109 promotes genome stability by acetylating histone H3 on lysine 56. Science. 2007;315: 649–652. doi:10.1126/science.1135862

20. Burgess RJ, Zhou H, Han J, Zhang Z. A role for Gcn5 in replication-coupled nucleosome assembly. Mol Cell. 2010;37: 469–480. doi:10.1016/j.molcel.2010.01.020

21. Trujillo KM, Osley MA. A role for H2B ubiquitylation in DNA replication. Mol Cell. 2012;48: 734–746. doi:10.1016/j.molcel.2012.09.019

22. Rizzardi LF, Dorn ES, Strahl BD, Cook JG. DNA replication origin function is promoted by H3K4 di-methylation in Saccharomyces cerevisiae. Genetics. 2012;192: 371–384. doi:10.1534/genetics.112.142349

23. Costas C, de la Paz Sanchez M, Stroud H, Yu Y, Oliveros JC, Feng S, et al. Genome-wide mapping of Arabidopsis thaliana origins of DNA replication and their associated epigenetic marks. Nat Struct Mol Biol. 2011;18: 395–400. doi:10.1038/nsmb.1988

24. Valenzuela MS, Chen Y, Davis S, Yang F, Walker RL, Bilke S, et al. Preferential localization of human origins of DNA replication at the 5’-ends of expressed genes and at evolutionarily conserved DNA sequences. PLoS ONE. 2011;6: e17308. doi:10.1371/journal.pone.0017308

25. Miotto B, Ji Z, Struhl K. Selectivity of ORC binding sites and the relation to replication timing, fragile sites, and deletions in cancers. Proc Natl Acad Sci USA. 2016;113: E4810–4819. doi:10.1073/pnas.1609060113

26. Rondinelli B, Schwerer H, Antonini E, Gaviraghi M, Lupi A, Frenquelli M, et al. H3K4me3 demethylation by the histone demethylase KDM5C/JARID1C promotes DNA replication origin firing. Nucleic Acids Res. 2015;43: 2560–2574. doi:10.1093/nar/gkv090

27. Lu F, Wu X, Yin F, Chia-Fang Lee C, Yu M, Mihaylov IS, et al. Regulation of DNA replication and chromosomal polyploidy by the MLL-WDR5-RBBP5 methyltransferases. Biol Open. 2016;5: 1449–1460. doi:10.1242/bio.019729

28. Beilharz TH, Harrison PF, Miles DM, See MM, Le UMM, Kalanon M, et al. Coordination of Cell Cycle Progression and Mitotic Spindle Assembly Involves Histone H3 Lysine 4 Methylation by Set1/COMPASS. Genetics. 2017;205: 185–199. doi:10.1534/genetics.116.194852

29. Sollier J, Lin W, Soustelle C, Suhre K, Nicolas A, Géli V, et al. Set1 is required for meiotic S-phase onset, double-strand break formation and middle gene expression. EMBO J. 2004;23: 1957–1967. doi:10.1038/sj.emboj.7600204

30. Sun Z-W, Allis CD. Ubiquitination of histone H2B regulates H3 methylation and gene silencing in yeast. Nature. 2002;418: 104–108. doi:10.1038/nature00883

31. Delamarre A, Barthe A, de la Roche Saint-André C, Luciano P, Forey R, Padioleau I, et al. MRX Increases Chromatin Accessibility at Stalled Replication Forks to Promote Nascent DNA Resection and Cohesin Loading. Mol Cell. 2020;77: 395–410.e3. doi:10.1016/j.molcel.2019.10.029

32. Chong SY, Cutler S, Lin J-J, Tsai C-H, Tsai H-K, Biggins S, et al. H3K4 methylation at active genes mitigates transcription-replication conflicts during replication stress. Nat Commun. 2020;11: 809. doi:10.1038/s41467-020-14595-4

33. Liang C, Weinreich M, Stillman B. ORC and Cdc6p interact and determine the frequency of initiation of DNA replication in the genome. Cell. 1995;81: 667–676. doi:10.1016/0092-8674(95)90528-6

34. Dehé P-M, Dichtl B, Schaft D, Roguev A, Pamblanco M, Lebrun R, et al. Protein interactions within the Set1 complex and their roles in the regulation of histone 3 lysine 4 methylation. J Biol Chem. 2006;281: 35404–35412. doi:10.1074/jbc.M603099200

35. Guillemette B, Drogaris P, Lin H-HS, Armstrong H, Hiragami-Hamada K, Imhof A, et al. H3 lysine 4 is acetylated at active gene promoters and is regulated by H3 lysine 4 methylation. PLoS Genet. 2011;7: e1001354. doi:10.1371/journal.pgen.1001354

36. Bian C, Xu C, Ruan J, Lee KK, Burke TL, Tempel W, et al. Sgf29 binds histone H3K4me2/3 and is required for SAGA complex recruitment and histone H3 acetylation. EMBO J. 2011;30: 2829–2842. doi:10.1038/emboj.2011.193

37. Zhong Y, Nellimoottil T, Peace JM, Knott SRV, Villwock SK, Yee JM, et al. The level of origin firing inversely affects the rate of replication fork progression. J Cell Biol. 2013;201: 373–383. doi:10.1083/jcb.201208060

38. Viggiani CJ, Aparicio OM. New vectors for simplified construction of BrdU-Incorporating strains of Saccharomyces cerevisiae. Yeast. 2006;23: 1045–1051. doi:10.1002/yea.1406

39. Dillin A, Rine J. Roles for ORC in M phase and S phase. Science. 1998;279: 1733–1737. doi:10.1126/science.279.5357.1733

40. Suter B, Tong A, Chang M, Yu L, Brown GW, Boone C, et al. The origin recognition complex links replication, sister chromatid cohesion and transcriptional silencing in Saccharomyces cerevisiae. Genetics. 2004;167: 579–591. doi:10.1534/genetics.103.024851

41. Shimada K, Gasser SM. The origin recognition complex functions in sister-chromatid cohesion in Saccharomyces cerevisiae. Cell. 2007;128: 85–99. doi:10.1016/j.cell.2006.11.045

42. Wang H, Liu D, Wang Y, Qin J, Elledge SJ. Pds1 phosphorylation in response to DNA damage is essential for its DNA damage checkpoint function. Genes Dev. 2001;15: 1361–1372. doi:10.1101/gad.893201

43. Kim EM, Burke DJ. DNA damage activates the SAC in an ATM/ATR-dependent manner, independently of the kinetochore. PLoS Genet. 2008;4: e1000015. doi:10.1371/journal.pgen.1000015

44. Zhao X, Muller EG, Rothstein R. A suppressor of two essential checkpoint genes identifies a novel protein that negatively affects dNTP pools. Mol Cell. 1998;2: 329–340. doi:10.1016/s1097-2765(00)80277-4

45. Petrini JH, Bressan DA, Yao MS. The RAD52 epistasis group in mammalian double strand break repair. Semin Immunol. 1997;9: 181–188. doi:10.1006/smim.1997.0067

46. Duch A, Felipe-Abrio I, Barroso S, Yaakov G, García-Rubio M, Aguilera A, et al. Coordinated control of replication and transcription by a SAPK protects genomic integrity. Nature. 2013;493: 116–119. doi:10.1038/nature11675

47. Zhao H, Zhu M, Limbo O, Russell P. RNase H eliminates R-loops that disrupt DNA replication but is nonessential for efficient DSB repair. EMBO Rep. 2018;19. doi:10.15252/embr.201745335

48. Matos DA, Zhang J-M, Ouyang J, Nguyen HD, Genois M-M, Zou L. ATR Protects the Genome against R Loops through a MUS81-Triggered Feedback Loop. Mol Cell. 2020;77: 514–527.e4. doi:10.1016/j.molcel.2019.10.010

49. Chappidi N, Nascakova Z, Boleslavska B, Zellweger R, Isik E, Andrs M, et al. Fork Cleavage-Religation Cycle and Active Transcription Mediate Replication Restart after Fork Stalling at Co-transcriptional R-Loops. Mol Cell. 2020;77: 528–541.e8. doi:10.1016/j.molcel.2019.10.026

50. Mejlvang J, Feng Y, Alabert C, Neelsen KJ, Jasencakova Z, Zhao X, et al. New histone supply regulates replication fork speed and PCNA unloading. J Cell Biol. 2014;204: 29–43. doi:10.1083/jcb.201305017

51. Weinberger M, Ramachandran L, Feng L, Sharma K, Sun X, Marchetti M, et al. Apoptosis in budding yeast caused by defects in initiation of DNA replication. J Cell Sci. 2005;118: 3543–3553. doi:10.1242/jcs.02477

52. Walter D, Matter A, Fahrenkrog B. Loss of histone H3 methylation at lysine 4 triggers apoptosis in Saccharomyces cerevisiae. PLoS Genet. 2014;10: e1004095. doi:10.1371/journal.pgen.1004095

53. Rowe LA, Degtyareva N, Doetsch PW. DNA damage-induced reactive oxygen species (ROS) stress response in Saccharomyces cerevisiae. Free Radic Biol Med. 2008;45: 1167–1177. doi:10.1016/j.freeradbiomed.2008.07.018

54. Ge XQ, Jackson DA, Blow JJ. Dormant origins licensed by excess Mcm2-7 are required for human cells to survive replicative stress. Genes Dev. 2007;21: 3331–3341. doi:10.1101/gad.457807

55. Ibarra A, Schwob E, Méndez J. Excess MCM proteins protect human cells from replicative stress by licensing backup origins of replication. Proc Natl Acad Sci USA. 2008;105: 8956–8961. doi:10.1073/pnas.0803978105

56. Poli J, Tsaponina O, Crabbé L, Keszthelyi A, Pantesco V, Chabes A, et al. dNTP pools determine fork progression and origin usage under replication stress. EMBO J. 2012;31: 883–894. doi:10.1038/emboj.2011.470

57. Bacal J, Moriel-Carretero M, Pardo B, Barthe A, Sharma S, Chabes A, et al. Mrc1 and Rad9 cooperate to regulate initiation and elongation of DNA replication in response to DNA damage. EMBO J. 2018;37. doi:10.15252/embj.201899319

58. Cimprich KA, Cortez D. ATR: an essential regulator of genome integrity. Nat Rev Mol Cell Biol. 2008;9: 616–627. doi:10.1038/nrm2450

59. Singh A, Xu Y-J. The Cell Killing Mechanisms of Hydroxyurea. Genes (Basel). 2016;7. doi:10.3390/genes7110099

60. Heichinger C, Penkett CJ, Bähler J, Nurse P. Genome-wide characterization of fission yeast DNA replication origins. EMBO J. 2006;25: 5171–5179. doi:10.1038/sj.emboj.7601390

61. Blitzblau HG, Chan CS, Hochwagen A, Bell SP. Separation of DNA replication from the assembly of break-competent meiotic chromosomes. PLoS Genet. 2012;8: e1002643. doi:10.1371/journal.pgen.1002643

62. Gibson DG, Bell SP, Aparicio OM. Cell cycle execution point analysis of ORC function and characterization of the checkpoint response to ORC inactivation in Saccharomyces cerevisiae. Genes Cells. 2006;11: 557–573. doi:10.1111/j.1365-2443.2006.00967.x

63. Shimada K, Pasero P, Gasser SM. ORC and the intra-S-phase checkpoint: a threshold regulates Rad53p activation in S phase. Genes Dev. 2002;16: 3236–3252. doi:10.1101/gad.239802

64. Maya-Mendoza A, Moudry P, Merchut-Maya JM, Lee M, Strauss R, Bartek J. High speed of fork progression induces DNA replication stress and genomic instability. Nature. 2018;559: 279–284. doi:10.1038/s41586-018-0261-5

65. Tsang CK, Liu Y, Thomas J, Zhang Y, Zheng XFS. Superoxide dismutase 1 acts as a nuclear transcription factor to regulate oxidative stress resistance. Nat Commun. 2014;5: 3446. doi:10.1038/ncomms4446

66. Acquaviva L, Székvölgyi L, Dichtl B, Dichtl BS, de La Roche Saint André C, Nicolas A, et al. The COMPASS subunit Spp1 links histone methylation to initiation of meiotic recombination. Science. 2013;339: 215–218. doi:10.1126/science.1225739

67. Schneider CA, Rasband WS, Eliceiri KW. NIH Image to ImageJ: 25 years of image analysis. Nat Methods. 2012;9: 671–675. doi:10.1038/nmeth.2089

68. Mann RK, Grunstein M. Histone H3 N-terminal mutations allow hyperactivation of the yeast GAL1 gene in vivo. EMBO J. 1992;11: 3297–3306.

